# Swr1 mediated H2A.Z^Pht1^ incorporation designates centromere DNA for *de novo* CENP-A^Cnp1^ assembly

**DOI:** 10.1101/215962

**Authors:** Raghavendran Kulasegaran-Shylini, Lakxmi Subramanian, Alastair R. W. Kerr, Christos Spanos, Juri Rappsilber, Robin C. Allshire

## Abstract

The underlying hallmark of centromeres is the presence of specialized nucleosomes in which histone H3 is replaced by CENP-A. The events that mediate the installation of CENP-A in place of H3 remain poorly characterized. H2A.Z is linked to transcriptional competence and associates with mammalian centromeres. We find that H2A.Z^Pht1^ and the Swr1 complex are enriched in fission yeast CENP-A^Cnp1^ chromatin. Our analysis shows that Swr1, Msc1 and H2A.Z^Pht1^ are required to maintain CENP-A^Cnp1^ chromatin integrity. Cell cycle analyses demonstrate that H2A.Z^Pht1^ is deposited in S phase, coincident with the deposition of placeholder H3, and prior to CENP-A^Cnp1^ replenishment in G2. Establishment assays reveal that H2A.Z^Pht1^ and Swr1 are required for *de novo* assembly of CENP-A^Cnp1^ onto naïve centromere DNA. We propose that features akin to promoters within centromere DNA program the incorporation of H2A.Z^Pht1^ via Swr1, and mediate the replacement of resident H3 nucleosomes with CENP-A nucleosomes thereby defining centromeres.

## INTRODUCTION

Centromeres are the locations on chromosomes where kinetochores, the protein machinery responsible for accurate chromosome segregation, assemble. Many eukaryotic centromeres are distinguished by the presence of repetitive elements such as alpha satellite (human), minor satellite (mouse), cen180 repeats (plants), and inner & outer centromere repeats (fission yeast – *S. pombe*) (McKinley and Cheeseman, 2016; Muller and Almouzni, 2017). An underlying hallmark of many active centromeres is the presence of specialized nucleosomes in which canonical histone H3 is replaced by the centromere-specific histone H3 variant CENP-A (also known as cenH3; CID in *Drosophila*, Cnp1 in *S. pombe*). In addition, CENP-A chromatin is often flanked by, or interspersed with, heterochromatin (McKinley and Cheeseman, 2016; Muller and Almouzni, 2017).

Artificial tethering of CENP-A, or its assembly factors, induces the assembly of CENP-A and kinetochores at non-centromeric locations (Barnhart et al., 2011; Barrey and Heun, 2017; Mendiburo et al., 2011); thus, CENP-A incorporation alone is sufficient to mediate kinetochore assembly on non-centromeric sequences. Deletion of endogenous centromere sequences can result in the assembly of CENP-A and kinetochores at unusual chromosomal locations resulting in neocentromere formation (Ishii et al., 2008; Ketel et al., 2009; Marshall et al., 2008). Such observations suggest that DNA sequence itself is not of prime importance in specifying where CENP-A chromatin is assembled. Consequently, once incorporated at a chromosomal location CENP-A chromatin and the associated kinetochore have an innate ability to persist at that location through multiple cell divisions via an epigenetic positive feedback mechanism that ensures CENP-A replenishment every cell cycle (Burrack et al., 2016; Mendiburo et al., 2011). Nevertheless, other analyses indicate that centromere DNA sequences themselves are important. The introduction of human, mouse or *S. pombe* centromeric DNA into their respective cells shows that such sequences are efficient and preferred substrates for *de novo* CENP-A chromatin and kinetochore assembly ( Barrey and Heun, 2017). Hence, specific features within these centromeric DNAs must ensure their recognition and program a series of events resulting in the installation of CENP-A, instead of histone H3 nucleosomes, on these sequences. In support of this view, the binding of CENP-B to satellite repeats has been shown to be required to direct efficient *de novo* CENP-A and kinetochore assembly in mammalian cells (Ohzeki et al., 2002; Okada et al., 2007). However, a paradox is apparent as the chromosomal locations where human neocentromeres form lack satellite DNA and CENP-B binding sites (du Sart et al., 1997; Marshall et al., 2008).

The specific replenishment of CENP-A at centromeres during the cell cycle is ensured by temporally separating its assembly into chromatin from that of bulk H3 chromatin during S phase. During human centromere replication parental CENP-A is distributed to both daughter strands thereby reducing its levels at each centromere. H3.3 appears to be incorporated as a placeholder during S phase and is replaced with new CENP-A in G1 (Dunleavy et al., 2011; Jansen et al., 2007). The timing of new *Drosophila* CENP-A^CID^ incorporation is cell type dependent occurring either during mitosis or late telophase/early G1 (Erhardt et al., 2008; Mellone et al., 2011; Schuh et al., 2007). Microscopic analyses of single *S. pombe* cells revealed that CENP-A^Cnp1^ levels at the centromere cluster decline in S phase and are replenished in G2 indicating that, as in metazoa, CENP-A^Cnp1^ deposition is uncoupled from replication (Lando et al., 2012). Recent chromatin immunoprecipitation (ChIP) analyses demonstrate that newly synthesized CENP-A^Cnp1^ is deposited at *S. pombe* centromeres during G2 where it replaces placeholder histone H3 incorporated during the preceding S phase (Shukla et al., 2017).

Centromere DNA is transcribed in yeast, plants, flies and mammals (Duda et al., 2017). It is known that transcription can mediate histone replacement suggesting that transcription-coupled processes may promote incorporation of CENP-A in place of histone H3. In *S. pombe*, flanking heterochromatin plays a key role in the *de novo* establishment of CENP-A^Cnp1^ chromatin (Folco et al., 2008; Kagansky et al., 2009). Thus, in addition to CENP-A specific factors, the surrounding chromatin context can be pivotal in promoting CENP-A deposition. Related to this, following deletion of an endogenous *S. pombe* centromere, neocentromeres can form close to telomeres (Ishii et al., 2008). A feature of some neocentromeres is their relatively low levels of H2A.Z^Pht1^ occupancy prior to CENP-A^Cnp1^ incorporation (Ogiyama et al., 2013). In contrast, H2A.Z is enriched at mammalian centromeres and is essential for normal chromosome segregation (Foltz et al., 2006; Greaves et al., 2007; Rangasamy et al., 2004). Together these findings suggest that multiple redundant features earmark centromeric DNA sequences as preferred sites for CENP-A and kinetochore assembly.

The three *S. pombe* centromeres (*cen1*, *cen2*, and *cen3*) consist of two distinct regions of chromatin: outer repeats (*otr/dg-dh*) coated in H3K9me-dependent heterochromatin flank the three central domains (*cnt1*, *cnt2* and *cnt3*; each ~10 kb) where CENP-A^Cnp1^ nucleosomes and kinetochores are normally assembled (Figure 1A). In establishment assays CENP-A^Cnp1^ cannot be assembled *de novo* on minichromosomes bearing central domain DNA alone, however, CENP-A^Cnp1^ over-expression permits its assembly in place of H3 on this DNA. Central domain chromatin exhibits pervasive low quality transcription with a high density of convergent and divergent promoters. CENP-A^Cnp1^ assembly on central domain DNA is stimulated by reduced RNAPII transcription-coupled H3 chromatin reassembly or increased RNAPII stalling (Catania et al., 2015; Choi et al., 2011; Choi et al., 2012). Such observations suggest that the central domains of *S. pombe* centromeres may be programmed to replace H3 with CENP-A^Cnp1^, specifically within these regions, through transcription-coupled mechanisms. A variety of processes also operate to prevent the accumulation of CENP-A at non-centromeric locations (Choi et al., 2012; Lacoste et al., 2014; Ohkuni et al., 2016; Ranjitkar et al., 2010), and a recent study has suggested that Ino80-mediated H3 eviction favors CENP-A^Cnp1^ chromatin assembly (Choi et al., 2017). Thus, distinct and overlapping processes converge to ensure CENP-A deposition at only one location per chromosome.

**Figure 1:**
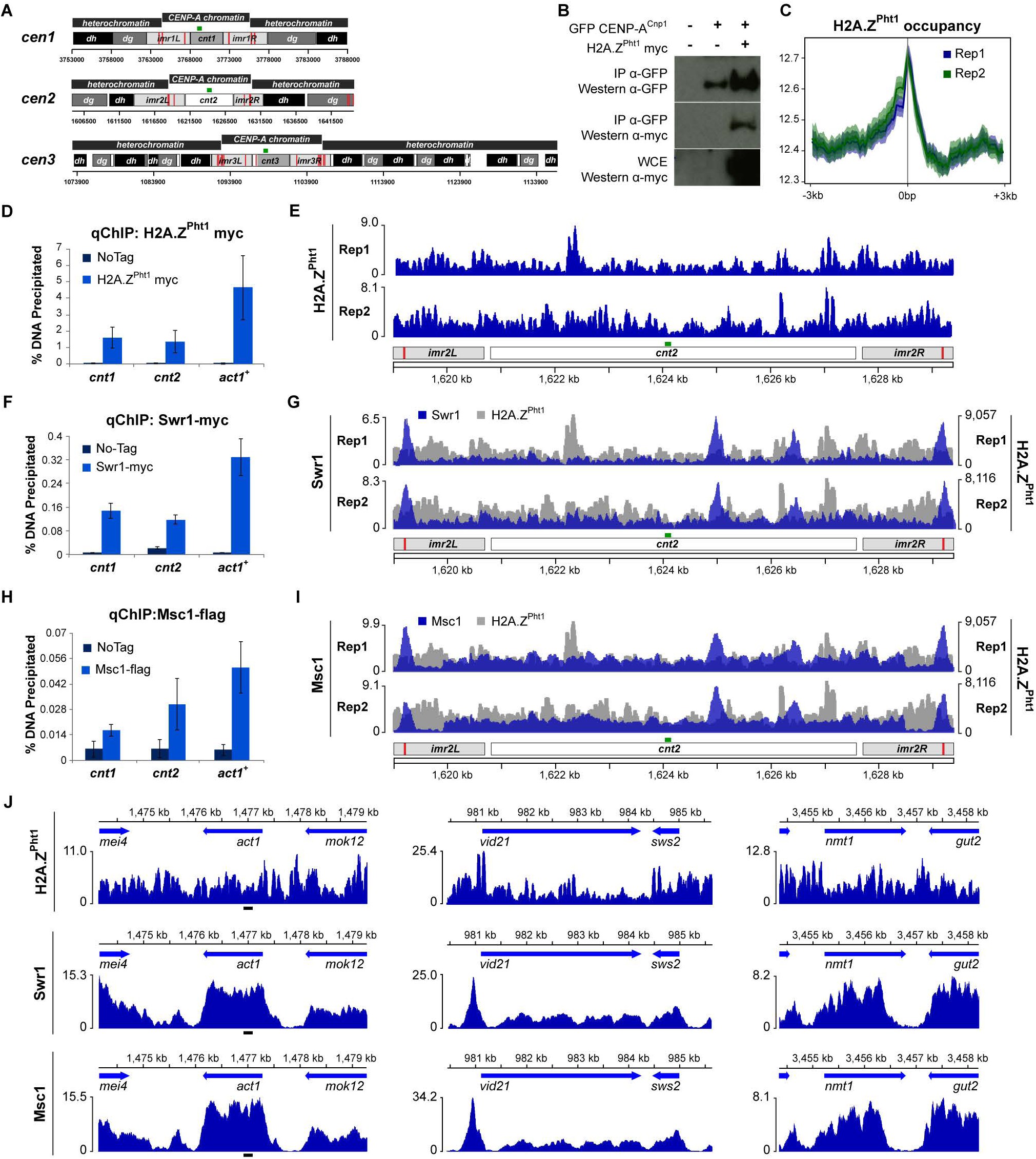
H2A.Z^Pht1^ and Swr1C associate with centromeric chromatin. (A) Schematic of *pombe* centromeres. CENP-A^Cnp1^ coats *cnt* (central core) and *imr* (inner repeats). Heterochromatin coats *dg*/*dh* outer repeats. Red lines: tRNA genes. Green lines: primer pairs for qChIP. (B) Western: GFP-CENP-A^Cnp1^ or H2A.Z^Pht1^–myc in immunoprecipitates (IP) or WCE as indicated. (C) H2A.Z^Pht1^-myc occupancy around TSS (± 3kb), two independent replicates. Scale: normalized RPKM. (D, E, F) qChIP enrichment of H2A.Z^Pht1^-myc, Swr1-myc and Msc1-FLAG at *cnt1*, *cnt2* or *act1*^*+*^ relative to no-tag control. Mean ± SEM of n≥3 independent experiments is shown. (E, G, I) ChIP-seq profile of H2A.Z^Pht1^, Swr1 and Msc1 enrichment over *cnt2* of *cen2*. For comparison, H2A.Z^Pht1^ profile is shown in grey (G an I). (J) ChIP-seq profiles of H2A.Z^Pht1^, Swr1 and Msc1 across select genes. Black line below act1: qChIP primer pair. Chromosome positions (kb; X axes) and normalized RPKM (Y axes) are as indicated (E-J). See also Figure S1, Table S1.

Here we identify H2A.Z^Pht1^ and components of the Swr1 complex (Swr1C) as being enriched in *S. pombe* CENP-A^Cnp1^ chromatin. Our analyses demonstrate that H2A.Z^Pht1^, Swr1 and Msc1 (a Swr1C associated protein) contribute to the maintenance of CENP-A^Cnp1^ at centromeres. H2A.Z^Pht1^ is transiently deposited at centromeres in S phase, coincident with H3 incorporation, but its levels subsequently decline along with H3 in advance of CENP-A^Cnp1^ incorporation in G2. Increased levels of H2A.Z^Pht1^ associate with centromere DNA lacking CENP-A^Cnp1^, and CENP-A^Cnp1^ levels are decreased at centromeres in the absence of H2A.Z^Pht1^ or Swr1. Remarkably, CENP-A^Cnp1^ chromatin, and thus functional kinetochores, cannot be established on centromeric DNA in the absence of H2A.Z^Pht1^ or Swr1. We propose that centromere DNA is configured to promote Swr1C-mediated H2A.Z incorporation, which is required for H3 to CENP-A^Cnp1^ exchange and CENP-A^Cnp1^ chromatin assembly.

## RESULTS

### H2A.Z^Pht1^ and Swr1C are enriched in CENP-A^Cnp1^ chromatin

To identify factors that associate with CENP-A^Cnp1^ chromatin and that might contribute to CENP-A^Cnp1^ deposition we immunoprecipitated GFP-CENP-A^Cnp1^ from cells expressing N-terminally GFP-tagged protein from its native promoter. Previous studies show GFP-CENP-A^Cnp1^ tagged cells to have essentially wild-type centromeric function (Lando et al., 2012). Following affinity selection GFP-CENP-A^Cnp1^ associated proteins were identified by LC-MS/MS (Table S1). The known CENP-A^Cnp1^ chromatin associated proteins Sim3^NASP^ and Cnp3^CENP-C^ were detected along with various chromatin remodeling factors including Hrp1^Chd1^, FACT and Ino80, known to affect CENP-A^Cnp1^ chromatin assembly in *S. pombe* (Choi et al., 2017; Choi et al., 2012; Walfridsson et al., 2005). Peptides corresponding to the RSC and SWI/SNF remodeling complexes were also detected along with subunits of the NuA4/Mst1^Kat5^ and Mst2^Kat7^ histone acetyltransferases. In addition, peptides for both the histone variant H2A.Z (encoded by the *pht1*^+^ gene in *S. pombe*) and shared components of the Swr1 and Ino80 complexes were detected. Swr1 is known to be required to replace H2A with H2A.Z (Kobor et al., 2004; Krogan et al., 2003; Mizuguchi et al., 2004) and in *S. pombe* H2A.Z^Pht1^ and Swr1C are required to maintain centromeric silencing and genome stability (Ahmed et al., 2007; Hou et al., 2010; Kim et al., 2009; Zofall et al., 2009). H2A.Z has previously been reported to be enriched in human CENP-A chromatin suggesting a possible role for H2A.Z in maintaining centromeric chromatin architecture (Foltz et al., 2006; Greaves et al., 2007; Nekrasov et al., 2012; Rangasamy et al., 2004).

Marker genes placed within the central domain of *S. pombe* centromeres (*cnt1:arg3*^*+*^) are transcriptionally repressed. Mutation of CENP-A^Cnp1^, or its loading factors Sim3^NASP^ or Scm3^HJURP^, cause reduced CENP-A^Cnp1^ incorporation within centromeric chromatin and alleviate this silencing (Dunleavy et al., 2007; Pidoux et al., 2009). Cells lacking genes encoding individual components of various chromatin-remodeling complexes associated with CENP-A^Cnp1^ chromatin were tested in centromere silencing assays. Deletion of Mst2^Kat7^, components of NuA4/Mst1^Kat5^ histone acetyltransferase (Eaf6 and Alp13), or non-essential components of the Ino80 complex (Arp5 and Arp8) failed to alleviate silencing of *cnt1:arg3*^*+*^ relative to controls, indicating that they do not appreciably affect CENP-A^Cnp1^ chromatin maintenance at centromeres. Cells lacking genes encoding H2A.Z^Pht1^ or components of the Swr1C (Swr1, Swc2, Swc3, Msc1 and Yaf9) exhibited partial alleviation of silencing (Figure S1A), suggesting that H2A.Z^Pht1^ and Swr1C might be involved in preserving CENP-A^Cnp1^ chromatin integrity.

Association of H2A.Z^Pht1^-myc with GFP-CENP-A^Cnp1^ was confirmed by co-immunoprecipitation from cell extracts (Figure 1B). Consistent with this, quantitative ChIP assays (qChIP) demonstrated H2A.Z^Pht1^-myc to be appreciably enriched within the central CENP-A^Cnp1^ chromatin domain (*cnt1* & *cnt2*) of centromeres (Figure 1D). Although the levels of H2A.Z^Pht1^-myc associated with the central domain is lower than that detected at the highly expressed *act1*^*+*^ gene, it is significantly enriched relative to an untagged control (Figure 1D). Previous analyses showed that H2A.Z^Pht1^ associates with the 5’ region of *S. pombe* genes (Buchanan et al., 2009; Hou et al., 2010; Zofall et al., 2009). Using ChIP-seq we also detected H2A.Z^Pht1^-myc close to the transcription start site (TSS) of genes (Figure 1C, J). In addition, we found H2A.Z^Pht1^-myc to be distributed throughout the central domains of all three centromeres, albeit at lower levels than promoter regions (Figure 1E, S1B). This detection of low H2A.Z^Pht1^-myc levels over the central domain differs from an earlier report suggesting H2A.Z^Pht1^ to be largely absent from the CENP-A^Cnp1^ chromatin region (Buchanan et al., 2009). ChIP-seq, as used here, generally provides greater sensitivity than ChIP-chip (Ho et al., 2011), thus the apparent discrepancy may reflect inherent differences between methodologies. We conclude that H2A.Z^Pht1^ associates with CENP-A^Cnp1^ and is present throughout the central domain of fission yeast centromeres.

### Swr1 and Msc1 are required for H2A.Z deposition within *S. pombe* centromeres

Swr1 is the main catalytic subunit of the Swr1C which mediates the incorporation of H2A.Z-H2B dimers replacing H2A-H2B dimers in nucleosomes, particularly at the +1 position adjacent to promoter associated nucleosome free regions (NFRs) (Ranjan et al., 2013; Yen et al., 2013). Msc1 is a KDM5-related protein that associates with *S. pombe* Swr1C but is absent in *S. cerevisiae*. Msc1 has also been implicated in H2A.Z^Pht1^ deposition and contributes to genome stability (Ahmed et al., 2007; Buchanan et al., 2009; Gao et al., 2017; Hou et al., 2010). Since H2A.Z^Pht1^-myc was measurably enriched within the central domain of centromere we next applied qChIP and ChIP-seq to determine if Swr1 and Msc1 also associate with the central domain. qChIP assays indicated that Swr1-myc and Msc1-FLAG are quantitatively enriched within the *cnt1* and *cnt2* regions of centromeres and on the *act1*^*+*^ gene relative to untagged controls (Figure 1F, H). ChIP-seq revealed that Swr1-myc and Msc1-FLAG generally exhibit greater enrichment over genic locations where H2A.Z^Pht1^ is deposited, such as the 5’ ends of genes (Figure 1J). In addition, both Swr1-myc and Msc1-FLAG were detected across the central domain of centromeres (Figure 1G, 1I, S1B). The association of Swr1 and Msc1 with the central domains suggests that they may mediate H2A.Z^Pht1^ incorporation within these regions.

Previous reports suggest that loss of Swr1 or Msc1 affected H2A.Z^Pht1^ deposition in euchromatin but resulted in either increased H2A.Z^Pht1^ deposition across centromeres or did not alter centromeric H2A.Z^Pht1^ occupancy (Buchanan et al., 2009; Hou et al., 2010). To determine if Swr1 and/or Msc1 are required for H2A.Z^Pht1^ deposition within central domains qChIP was used to assess the levels of H2A.Z^Pht1^-myc associated with *cnt1*, *cnt2* and the *act1*^*+*^ gene in wild-type cells (*wt*) and cells lacking Swr1 (*swr1∆*) or Msc1 (*msc1∆*). H2A.Z^Pht1^-myc levels were noticeably reduced relative to wild-type at all three locations in the absence of Swr1 or Msc1 (Figure 2A). ChIP-seq demonstrated that H2A.Z^Pht1^-myc levels at the 5’ ends of a cohort of genes is greatly reduced in *swr1∆* and *msc1∆* cells compared to wild-type, confirming that Swr1 and Msc1 are indeed required for H2A.Z^Pht1^ deposition in promoter-proximal NFRs (Figure 2B, 2C, S2). ChIP-seq analyses of H2A.Z^Pht1^-myc association with centromere regions demonstrated that both Swr1 and Msc1 are required for full incorporation of H2A.Z^Pht1^ throughout the central domain (Figure 2D). We conclude that both Swr1 and Msc1 are enriched across the central domain of centromeres where, as at the 5’ ends of many genes, they mediate H2A.Z^Pht1^ deposition.

**Figure 2:**
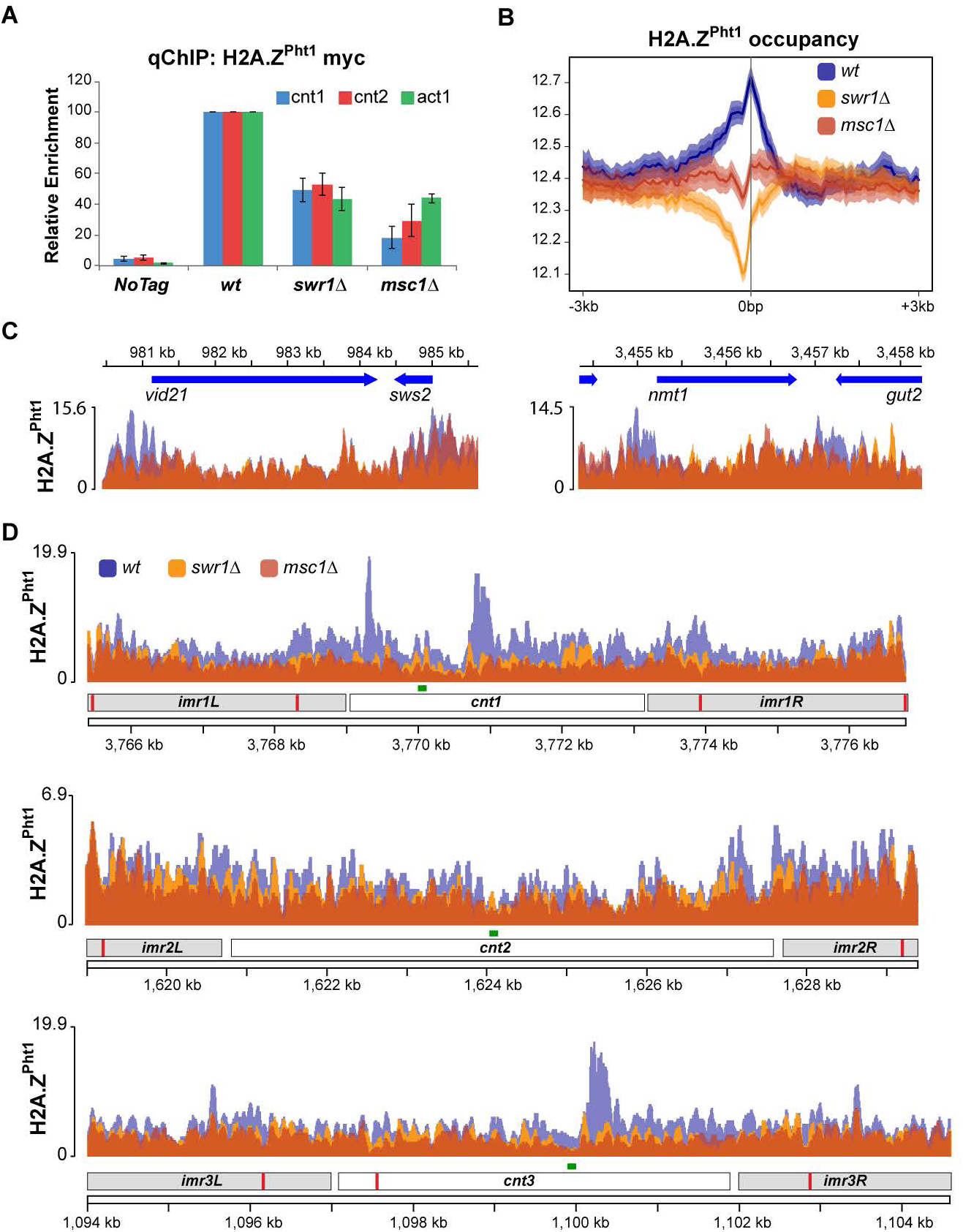
Swr1 and Msc1 are required for H2A.Z^Pht1^ deposition. (A) qChIP of H2A.Z^Pht1^ levels at *cnt1*, *cnt2* or *act1*^*+*^ in *swr1*∆ and *msc1*∆ cells normalized to wild-type (*wt*). Mean ± SEM of n≥3 independent experiments is shown. (B) H2A.Z^Pht1^-myc occupancy around TSS (± 3kb), in *wt*, *swr1∆* and *msc1∆*. Scale: normalized RPKM. (C) ChIP-seq profile for H2A.Z^Pht1^ across select genes in *wt*, *swr1∆* and *msc1∆*. Chromosome positions (kb) and annotation indicated. Scale: normalized RPKM. (D) ChIP-seq profiles of H2A.Z^Pht1^ enrichments over central domains of *cen1*, *2*, *3*. Green lines: *cnt1*, *2*, 3 primer pairs for qChIP. Chromosome positions (kb) and annotation indicated. Scale: normalized RPKM (C, D). See also Figure S2.

### H2A.Z^Pht1^ deposition within centromeres contributes to CENP-A^Cnp1^ maintenance

The partial alleviation of central domain reporter gene silencing in cells devoid of H2A.Z^Pht1^, Swr1C subunits or Msc1 (Figure S1A) suggests that reduced H2A.Z^Pht1^ deposition might affect CENP-A^Cnp1^ chromatin assembly. qChIP was therefore performed to assess the levels of CENP-A^Cnp1^ retained within centromeres in cells lacking H2A.Z^Pht1^, Swr1, or Msc1. CENP-A^Cnp1^ levels at both *cnt1* and *cnt2* were noticeably and reproducibly reduced in the absence of either H2A.Z^Pht1^, Swr1 or Msc1 (Figure 3A, 3C). ChIP-seq confirmed that CENP-A^Cnp1^ enrichment across the unique 7kb of the *cnt2* central domain, and other centromeres (*cnt1, cnt3*), was noticeably reduced in cells devoid of H2A.Z^Pht1^, Swr1, or Msc1 relative to wild type (Figure 3B, 3D, S3A, S3B). Thus, the deposition of H2A.Z^Pht1^ by Swr1C within the central domain of *S. pombe* centromeres appears to be important for the maintenance of wild-type CENP-A^Cnp1^ levels throughout the entire domain.

**Figure 3:**
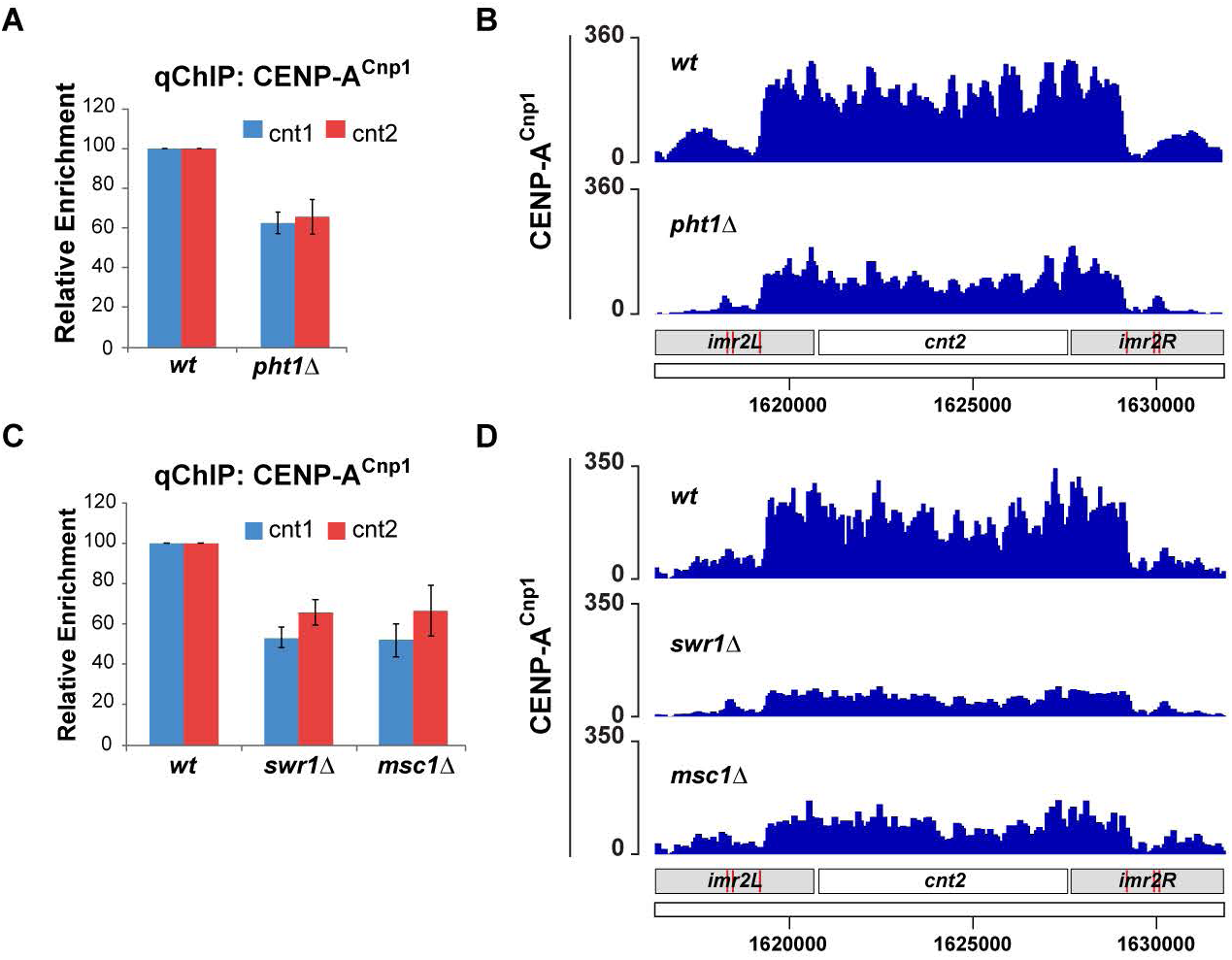
H2A.Z^Pht1^ and Swr1C affect CENP-A^Cnp1^ maintenance. (A & C) qChIP of CENP-A^Cnp1^ levels at *cnt1* or *cnt2* in *pht1∆* (A) or *swr1∆* and *msc1∆* (C) cells normalized to *wt*. Mean ± SEM of n≥3 independent experiments is shown. (B & D) CENP-A^Cnp1^ ChIP-seq profile over *cnt2* in *wt* and *pht1∆* (B) or *wt, swr1∆* and *msc1∆* (D). Chromosome positions (kb) and annotation indicated. Scale: normalized RPKM (C, D). See also Figure S3.

### H2A.Z^Pht1^ is enriched on naïve centromere DNA assembled in H3 chromatin

To test if H2A.Z^Pht1^ is prevalent when central domain DNA is assembled in H3 chromatin rather than CENP-A^Cnp1^ chromatin, minichromosomes (Figure 4A) harboring *cnt2* central domain DNA from *cen2* flanked by heterochromatin (pHCC2), or *cnt2* central domain DNA alone (pCC2), were transformed into wild-type cells where the central domain of endogenous *cen2* had been replaced with the central domain of *cen1* (*cc2∆::cc1*) (Figure 4B) (Catania et al., 2015). Following transformation, mitotic stability assays have previously shown that pHCC2 DNA directs efficient *de novo* CENP-A^Cnp1^ and functional centromere assembly. However, in the absence of a flanking heterochromatin repeat only H3 chromatin is assembled over the central domain of pCC2 (Figure 4C) (Catania et al., 2015; Folco et al., 2008). The chromatin assembled across *cnt2* DNA carried by pCC2 thus represents naïve central domain chromatin prior to its conversion to CENP-A^Cnp1^ chromatin. qChIP analyses confirmed that CENP-A^Cnp1^ chromatin was established on pHCC2 while pCC2 remained assembled in H3 chromatin (Figure 4D, E). Notably, higher levels of H2A.Z^Pht1^ were incorporated on *cnt2* when assembled in H3 chromatin on pCC2 as opposed to CENP-A^Cnp1^ chromatin on pHCC2 (Figure 4F). H2A.Z^Pht1^ incorporation on *act1*^+^ was unaltered by the presence of these minichromosomes (Figure S4A). Thus H2A.Z^Pht1^ is more highly enriched in central domain chromatin when assembled in naïve H3 chromatin, prior to or without CENP-A^Cnp1^ assembly. The incorporation of this higher level of H2A.Z^Pht1^ on pCC2 is also Swr1 and Msc1 dependent (Figure 4H). Since less H3 is deposited when H2A.Z^Pht1^ is present the incorporation of H2A.Z^Pht1^ by Swr1 within central domains may be required to destabilize H3 nucleosomes to elicit CENP-A^Cnp1^ deposition.

**Figure 4:**
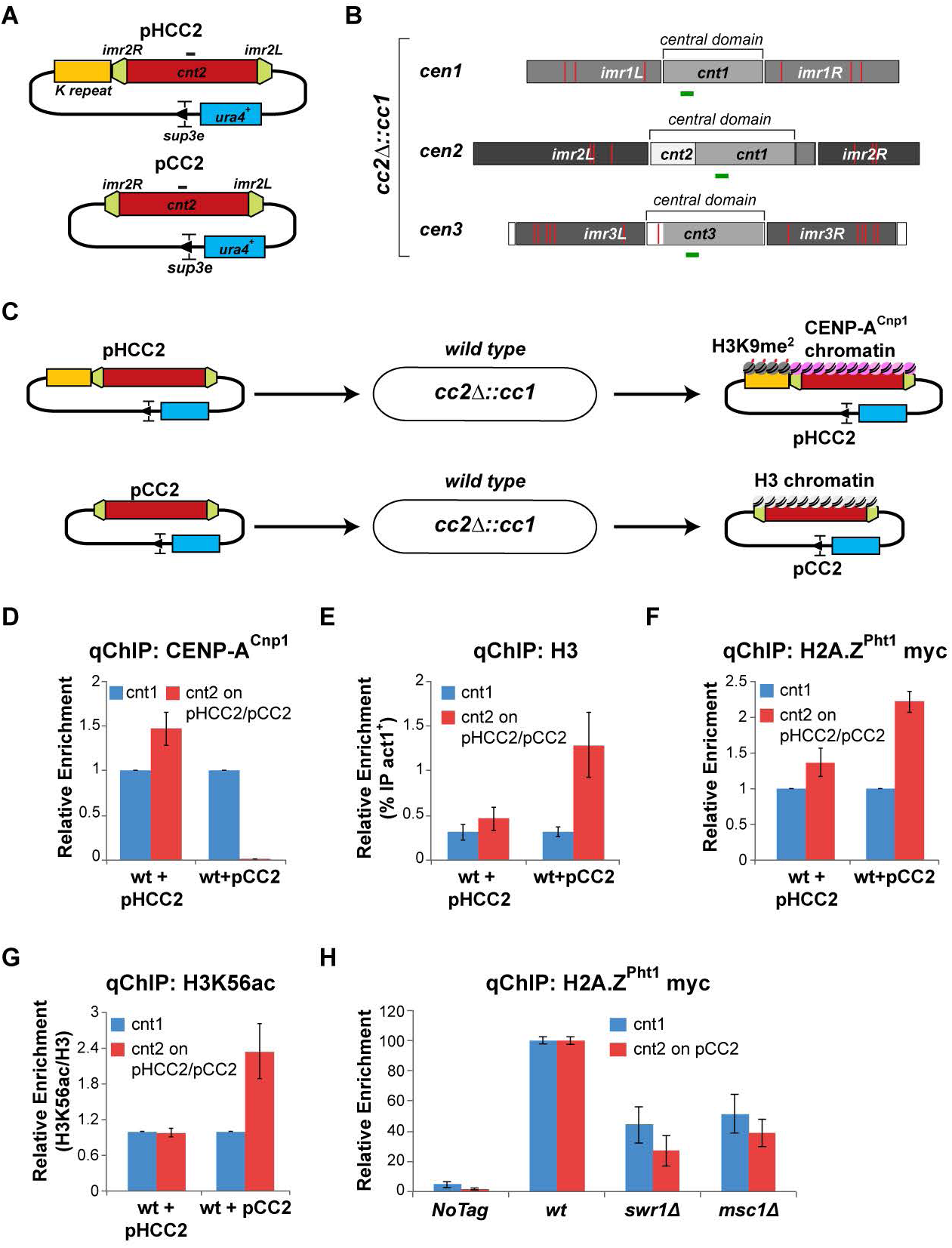
H2A.Z^Pht1^ is assembled on naïve centromeric DNA. (A) Schematic of minichromosomes carrying *cnt2* with flanking heterochromatin (pHCC2) or *cnt2* alone (pCC2). Black line: primer pair for pHCC2 of pHcc2 specific qChIP. (B) minichromosome recipient cells contain *cc2∆::cc1* where part of *cnt2* is replaced by 5.5 kb of cnt1 DNA. Green lines: primer pairs for *cnt1* and *cnt3* qChIP. (C) Schematic of establishment assay and expected outcomes following transformation of *wild-type cc2∆::cc1* cells with pHCC2 or pCC2. (D-G) qChIP of CENP-A^Cnp1^ (D), H3 (E) H2A.Z^Pht1^-myc (F) H3K56ac (H) levels at *cnt2* unique to pHCC2 or pCC2 minichromosomes (normalized to *cnt1*). Mean ± SEM of n≥3 independent experiments is shown. (H) qChIP of H2A.Z^Pht1^ levels at endogenous *cnt1* or *cnt2*, carried by pHCC2 or pCC2, in *swr1∆* and *msc1∆* cells normalized to *wt*. Mean ± SEM of n≥3 independent experiments is shown. See also Figure S4

The central domains of *S. pombe* centromeres are peppered with numerous transcriptional start sites and promoters, which direct pervasive low quality transcription (Catania et al., 2015; Choi et al., 2011). H3K56 acetylation is associated with freshly deposited histone H3 and is enriched in the dynamic or ‘hot’ H2A.Z^Pht1^ containing nucleosomes that reside at the +1 position downstream of many RNAPII promoters and throughout the bodies of highly transcribed genes (Marquardt et al., 2014; Rege et al., 2015). In agreement with this, high H3K56ac levels were detected on the highly expressed *act1*^+^ gene (Fig S4B). Higher levels of H3K56ac relative to total H3 were evident on the central *cnt2* domain of pCC2 compared to that of pHCC2, or at endogenous *cnt1* assembled in CENP-A^Cnp1^ (Figure 4G). The presence of high levels of both H2A.Z^Pht1^ and H3K56ac on pCC2*-cnt2* suggests that central domain DNA directs the assembly of unstable, ‘hot’ H3 nucleosomes, which likely designate these regions for CENP-A^Cnp1^ incorporation.

### H2A.Z^Pht1^ deposition at centromeres is cell cycle regulated

During replication, histone H3 is deposited as a placeholder prior to the deposition of new CENP-A^Cnp1^ in G2 when it is replenished to maximum levels (Shukla et al., 2017). To determine if the levels of H2A.Z^Pht1^ on the central domain at centromeres fluctuate during the cell cycle, relative to that of CENP-A^Cnp1^ and histone H3, cells were synchronized using the well characterized *cdc25-22* temperature sensitive mutation (Moreno et al., 1989). G2 blocked *cdc25-22* cells were released into the cell cycle and samples collected at regular intervals (Figure 5A). Synchrony was monitored by measuring the proportion of cells undergoing cytokinesis (% septated cells), which in *S. pombe* is coincident with S phase (Kim and Huberman, 2001) (Figure 5B, S5A). qChIP showed that CENP-A^Cnp1^ levels decline at centromeres to a minimum during S phase and increase again to full levels during the subsequent G2 phase (Figure 5C, S5B). In contrast total H3 levels on the central domain increased during S phase and declined at G2 onset (Figure 5D, S5C). H2A.Z^Pht1^ levels mirrored that of H3, increasing during S phase and declining in advance of CENP-A^Cnp1^ deposition during G2 (Figure 5E, S5D). The relative levels of H3K56ac show a similar pattern consistent with the deposition of H3 as a placeholder during S phase (Figure 5F, S5E). H2A.Z^Pht1^ incorporation signifies the presence of dynamic nucleosomes that are likely to be unstable (Jin and Felsenfeld, 2007). The coincident rise and fall of H2A.Z^Pht1^, H3 and H3K56ac levels on the central domain of centromeres during S phase suggests that the incorporation of H2A.Z^Pht1^ into central domain chromatin may be required to remove placeholder H3 to allow subsequent deposition of new CENP-A^Cnp1^.

**Figure 5:**
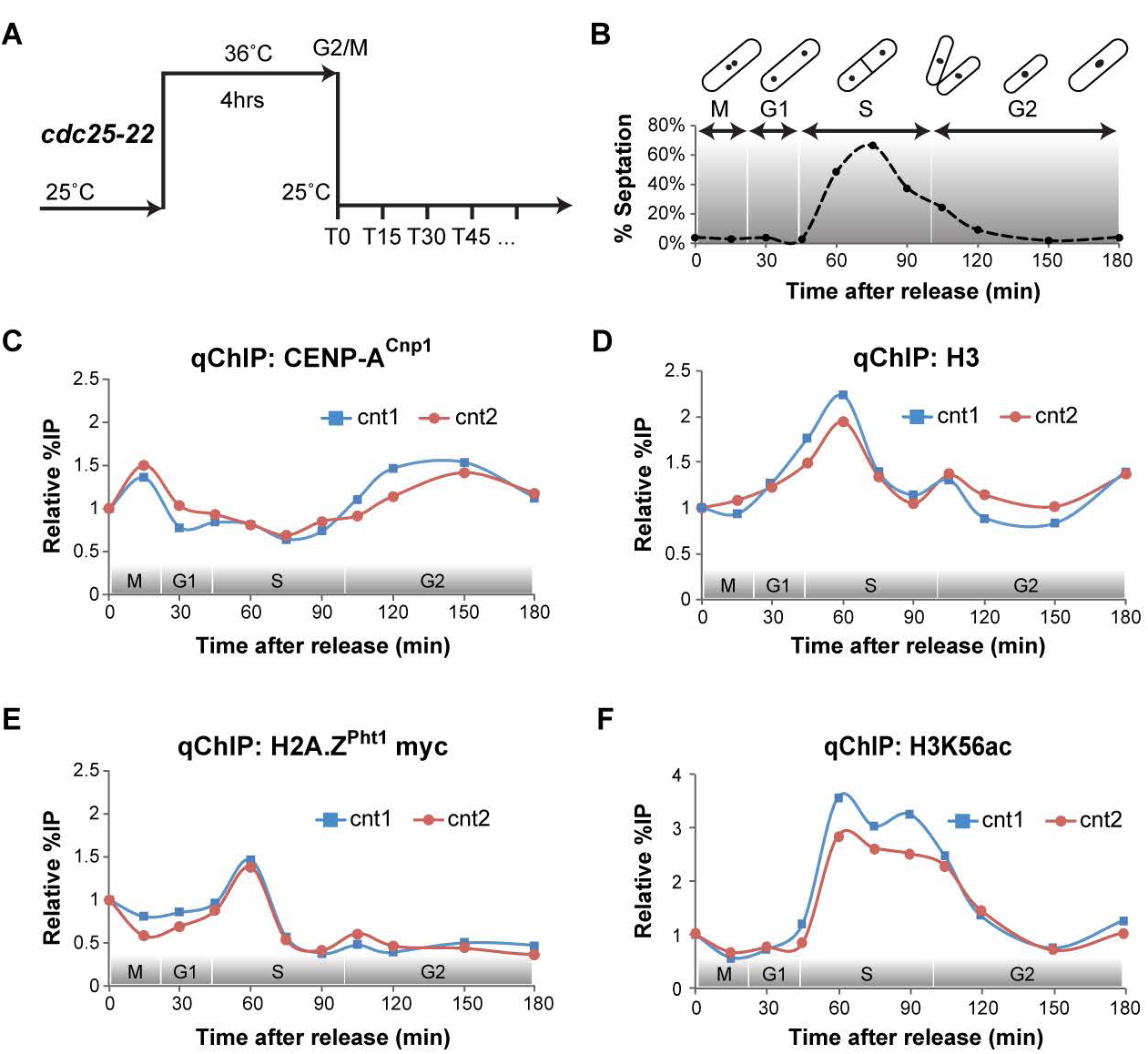
Cell cycle coordinated deposition of H2A.Z^Pht1^ with H3 precedes CENP-A^Cnp1^ replenishment. (A) Scheme to generate G2 synchronized cultures expressing H2A.Z^Pht1^-myc with a *cdc2-22* cell cycle block-release. Samples were harvested at indicated times (B-F). (B) Septation index plot to monitor cell cycle synchrony. Expected cell cycle phases shown above. qChIP for (C) CENP-A^Cnp1^, (D) H3, (E) H2A.Z^Pht1^ and (F) H3K56ac levels at *cnt1* and *cnt2* through cell cycle. See independent duplicate experiment in Figure S5.

### H2A.Z^Pht1^ and Swr1 are required to establish CENP-A^Cnp1^ chromatin and functional centromeres

The reduced levels of CENP-A^Cnp1^ associated with centromeres in cells lacking H2A.Z^Pht1^, Swr1 or Msc1 indicate that H2A.Z^Pht1^ is required to maintain CENP-A^Cnp1^ chromatin at centromeres. To determine if H2A.Z^Pht1^, Swr1 or Msc1 are required for the *de novo* establishment of CENP-A^Cnp1^ chromatin, the pHCC2 minichromosome was transformed into wild-type, *clr4∆*, *pht1∆*, *swr1∆,* or *msc1∆* cells. CENP-A^Cnp1^ establishment assays show that while pHCC2 DNA was able to direct efficient *de novo* establishment of functional centromeres in wild-type cells, little or no CENP-A^Cnp1^ was assembled on pHCC2 introduced into *pht1∆* or *swr1∆* cells, similar to *clr4∆* cells that lack heterochromatin (Figure 6A). qChIP revealed that as with *clr4∆* cells, no CENP-A^Cnp1^ was assembled on pHCC2 (*cnt2*) in *pht1∆* or *swr1∆* cells whereas only low levels were detected on pHCC2 in *msc1∆* cells (Figure 6B). Reciprocally, high levels of H3 remained in place on pHCC2 in the absence of CENP-A^Cnp1^ assembly in *pht1∆* or *swr1∆* cells (Figure 6C). The detection of higher H3 levels on pHCC2 in *pht1∆* and *swr1∆* compared to *clr4∆* cells is consistent with H2A.Z^Pht1^/Swr1 still operating to destabilize H3 nucleosomes on the central domain in *clr4∆* cells. In contrast, Msc1 is required for the maintenance of full CENP-A^Cnp1^ chromatin levels, but not establishment, thus reduced CENP-A^Cnp1^ is seen both at endogenous centromeres and on freshly introduced pHCC2.

**Figure 6:**
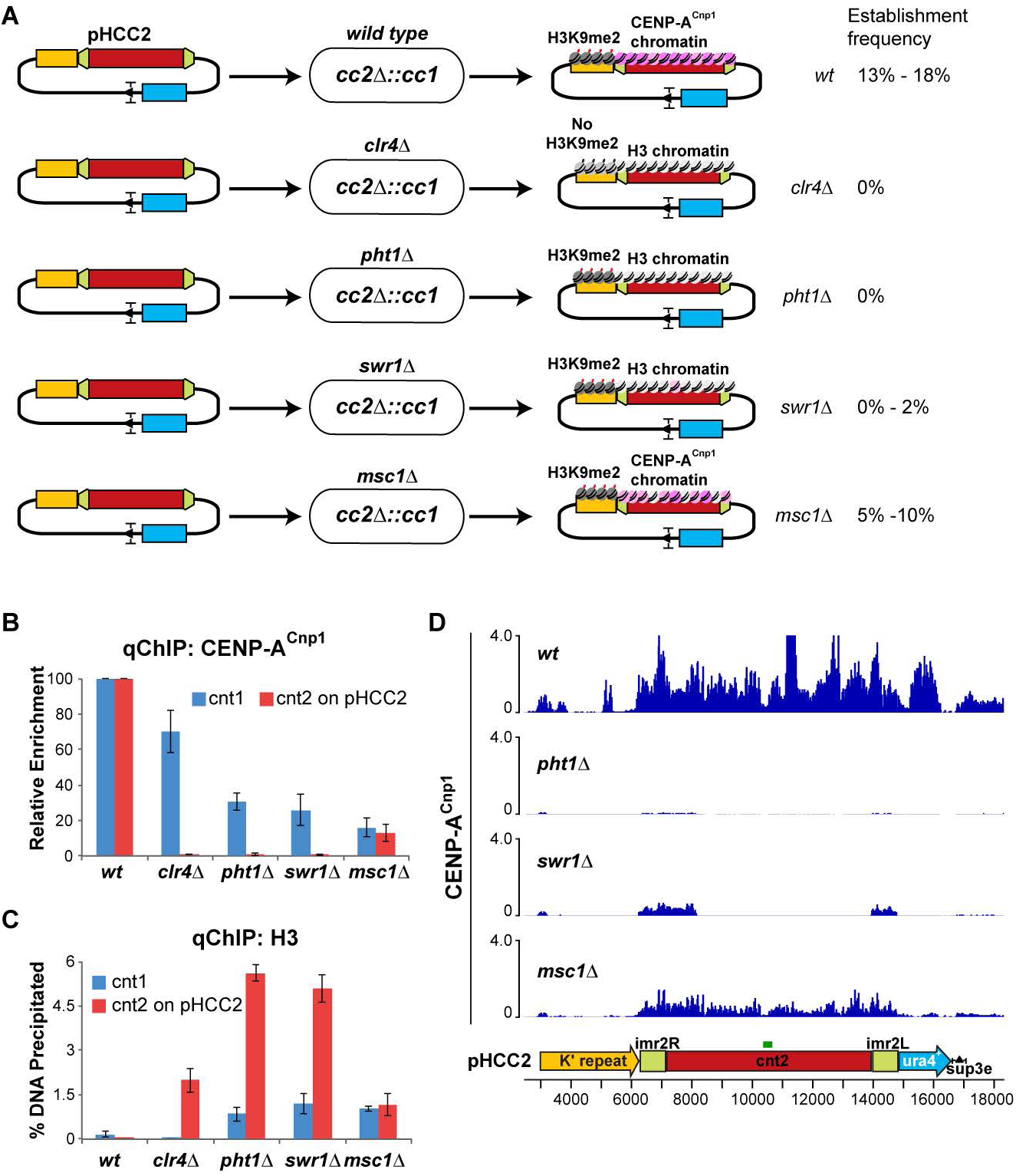
H2A.Z^Pht1^ and Swr1 are required for *de novo* CENP-A^Cnp1^ establishment. (A) Schematic of establishment assays using pHCC2 minichromosome in *cc2∆::cc1* cells in wild-type or mutant cells and observed outcome. (B & C) qChIP for CENP-A^Cnp1^ (B) or H3 (C) levels at endogenous *cnt1*, and *cnt2* on pHCC2 in *pht1∆*, *swr1∆* and *msc1∆* cells normalized to *wt*. Mean ± SEM of n≥3 independent experiments is shown. (D) ChIP-seq profiles for CENP-A^Cnp1^ across cnt2 of pHCC2 after transformation of wild-type, *pht1∆*, *swr1∆* or *msc1∆* cells. Scale represents normalized reads in RPKM. See also Figure S6.

ChIP-seq confirmed that no CENP-A^Cnp1^ chromatin was established across *cnt2* on the pHCC2 mini-chromosome following its transformation into cells lacking H2A.Z^Pht1^ or Swr1, while low levels of CENP-A^Cnp1^ were detectable in the absence of Msc1 (Figure 6D). The inability to establish CENP-A^Cnp1^ chromatin was not due to heterochromatin spreading across the central *cnt2* domain of pHCC2 since the levels of H3K9me2, the main heterochromatin mark, remained low over *cnt2* of pHCC2 relative to the outer *dg* repeat at endogenous centromeres that is assembled in H3K9me2-dependent heterochromatin (Figure S6A-C). We conclude that deposition of H2A.Z^Pht1^ within the central domain by Swr1 is critical for the *de novo* assembly of CENP-A^Cnp1^ chromatin at centromeres. In the absence of H2A.Z^Pht1^ deposition in central domains, resident H3 nucleosomes are more stable and consequently resist disassembly and replacement by CENP-A^Cnp1^ containing nucleosomes.

Acetylation of H2A.Z in its N-terminal region is associated with promoter-proximal nucleosomes of active *S. cerevisiae* genes where it is thought to promote transcription by regulating H2A.Z dynamics (Eirin-Lopez and Ausio, 2007; Millar et al., 2006). In *S. pombe,* H2A.Z^Pht1^ is similarly acetylated on four N-terminal lysines (K5, 7, 12 and 16) and this has been implicated in chromosome stability (Kim et al., 2009) and thus might affect *de novo* CENP-A^Cnp1^ chromatin assembly. qChIP analyses showed that H2A.Z^Pht1^ mutants mimicking the fully acetylated (4K→Q) or unacetylated state (4K→R) or lacking the N-terminus (cannot be acetylated) had essentially no defect in CENP-A^Cnp1^ maintenance at endogenous centromeres or on the levels of H2A.Z^Pht1^ associated with the central domain regions or *act1*^*+*^ (Figure S6D, E). In contrast however, only low levels of CENP-A^Cnp1^ were established on pHCC2 introduced into *pht1-∆N* and *pht1-4KR* mutants whereas CENP-A^Cnp1^ establishment was largely unaffected in *pht1-4KQ* cells (Figure S6F). This suggests that H2A.Z^Pht1^ dynamics within naïve central domain chromatin require N-terminal acetylation of H2A.Z^Pht1^, and its absence interferes with the mechanism that designates centromeres for CENP-A^Cnp1^ incorporation.

## DISCUSSION

Here we show that H2A.Z^Pht1^ is present throughout the central CENP-A^Cnp1^ domain of *S. pombe* centromeres. We find that Swr1C is required for H2A.Z^Pht1^ deposition in this region and that complete CENP-A^Cnp1^ chromatin integrity requires H2A.Z^Pht1^ and Swr1C. During the cell cycle H2A.Z^Pht1^ levels at centromeres increase during S phase, coincident with the deposition of placeholder H3 that is removed prior to CENP-A^Cnp1^ replenishment in G2. Using naïve centromere DNA templates we demonstrate that H2A.Z^Pht1^ and Swr1 are required to direct the *de novo* assembly of CENP-A^Cnp1^ chromatin on central domain DNA. Together our findings suggest a model where the deposition of H2A.Z^Pht1^ by Swr1 destabilizes resident H3 nucleosomes on centromere DNA promoting their replacement with CENP-A^Cnp1^ nucleosomes (Figure 7).

**Figure 7:**
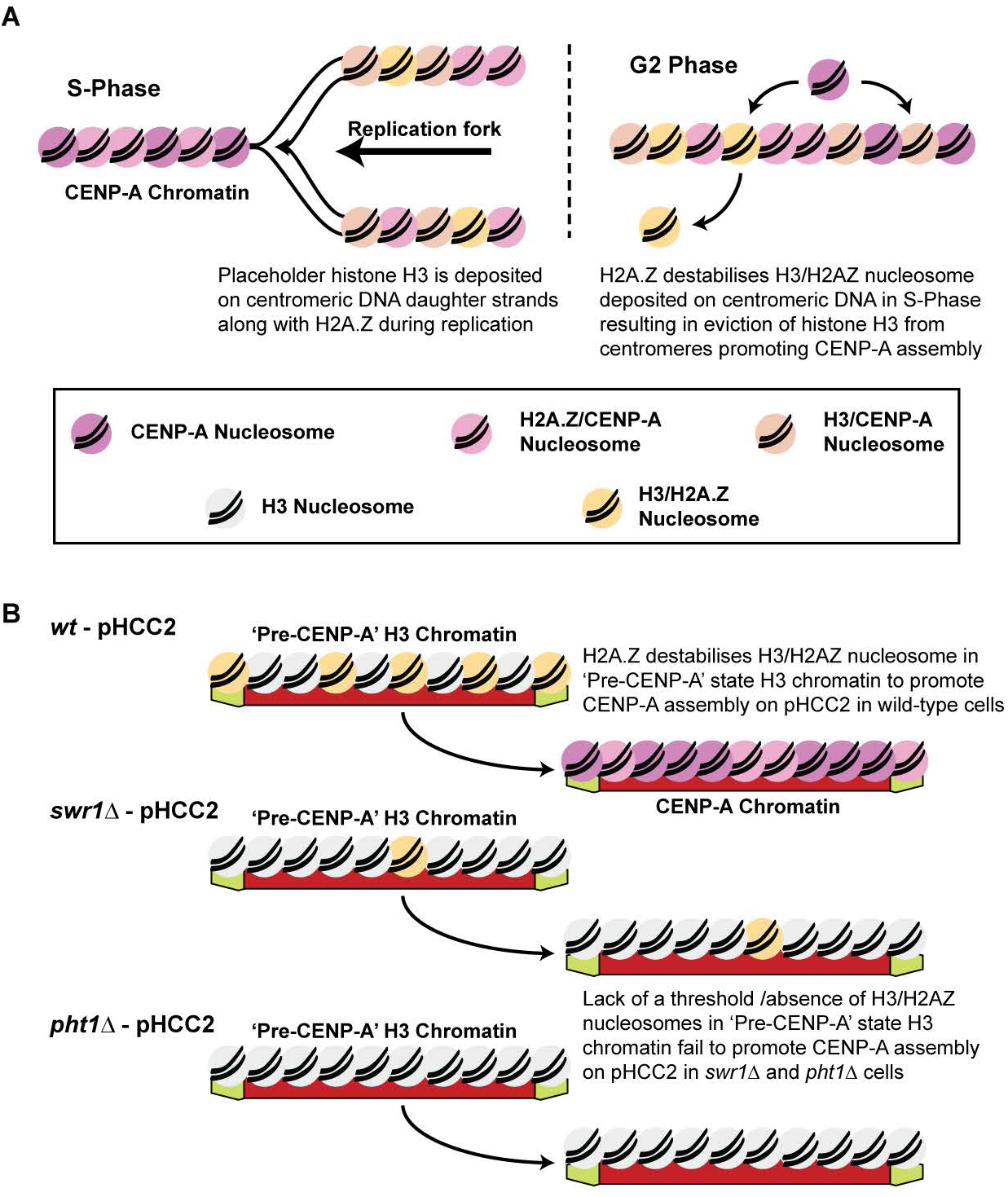
H2A.Z^Pht1^ and Swr1 mediated H3 eviction on centromeric DNA sequence promotes CENP-A^Cnp1^ assembly. (A) Model for CENP-A^Cnp1^ assembly through the cell cycle. As the fork progresses during S-phase parental CENP-A^Cnp1^ nucleosomes are distributed to sister chromatids. H2A.Z^Pht1^ containing H3 nucleosomes are deposited by dedicated chaperones to fill the gaps. These H2A.Z^Pht1^/H3 nucleosomes unstable and rapidly evicted from centromeres to enable CENP-A^Cnp1^ replenishment during G2. (B) Minichromosome pHCC2 minichromosomes may initially assemble dynamic H3 chromatin enriched for H2A.Z^Pht1^ (‘Pre-CENP-A’ state). H2A.Z^Pht1^ may then promote CENP-A^Cnp1^ assembly by ensuring H2A.Z/H3 nucleosomes disassembly. Little or no H2A.Z within central domain H3 chromatin on pHCC2 in *swr1∆* and *pht1∆* cells prevents conversion of ‘Pre-CENP-A’ H3 chromatin to CENP-A chromatin.

H2A.Z is enriched in promoter-proximal nucleosomes in many eukaryotes where it contributes to transcriptional competence (Albert et al., 2007; Raisner et al., 2005). Swr1C associates with yeast promoters via their NFRs and is required for the stepwise replacement of H2A-H2B with H2A.Z-H2B dimers in adjacent dynamic +1 nucleosomes (Ranjan et al., 2013; Yen et al., 2013). The dynamic nature of +1 nucleosomes is also supported by the presence of H3K56ac, a signature of newly deposited nucleosomes (Marquardt et al., 2014; Rege et al., 2015). Previous analyses showed that multiple transcriptional start sites are encoded by the central domains of *S. pombe* centromeres, and when not assembled in CENP-A^Cnp1^ chromatin they exhibit intrinsically low H3 nucleosome occupancy with high turnover (Catania et al., 2015; Choi et al., 2011; Shukla et al., 2017). Thus, central domains may essentially exhibit the properties equivalent to a large, or multiple tandem, NFRs. Consistent with this possibility we find that H2A.Z^Pht1^ occupancy is relatively low at endogenous centromeres compared to genic regions. The observed increase in H3 levels on naïve central domain DNA (pCC2 which lacks CENP-A^Cnp1^) in the absence of H2A.Z^Pht1^ or Swr1 (Figure 6C) also supports a role for H2A.Z^Pht1^ and Swr1 in destabilizing resident H3 nucleosomes. The coincident increase in H3, H3K56ac and H2A.Z^Pht1^, while CENP-A^Cnp1^ declines, during S phase is consistent with the deposition of H3 placeholder nucleosomes that contain H2A.Z^Pht1^. Subsequently, in G2, H3 and H3K56ac levels decline as CENP-A^Cnp1^ association rises. The relative timing of these events suggests that the incorporation of H2A.Z^Pht1^ by Swr1C might promote the replacement of H3 nucleosomes with CENP-A^Cnp1^ during the cell cycle. In support of this possibility, we find that CENP-A^Cnp1^ chromatin cannot be established on mini-chromosomes in cells lacking H2A.Z^Pht1^ or Swr1 whereas H3 is efficiently replaced with CENP-A^Cnp1^ chromatin in wild-type cells. Such establishment assays are extreme in that they test for the ability to assemble CENP-A^Cnp1^ rather than H3 chromatin *de novo* on a naïve substrate. Nevertheless, it is likely that similar mechanisms are involved in replacing placeholder H3 nucleosomes during the cell cycle to maintain CENP-A^Cnp1^ chromatin at established endogenous centromeres.

We envisage that the CENP-A^Cnp1^ chromatin domain is defined by numerous weak promoters scattered across the central domain and that these drive the formation of multiple +1-like dynamic nucleosomes where Swr1 mediates H2A/H2B-to-H2A.Z^Pht1^/H2B exchange, resulting in their destabilization. Loss of H2A.Z^Pht1^ or Swr1 prevents H2A.Z^Pht1^ deposition, thereby stabilizing H3 nucleosomes and preventing *de novo* CENP-A^Cnp1^ nucleosome assembly (Figure 7). At endogenous centromeres, where CENP-A^Cnp1^ chromatin and kinetochores are already assembled, additional mechanisms involving conserved CENP-A^Cnp1^-specific loading factors (i.e. Mis18, Scm3^HJURP^) override the loss of H2A.Z^Pht1^ and Swr1, allowing CENP-A^Cnp1^ maintenance, albeit at reduced levels. The overall density of unstable H2A.Z^Pht1^ containing H3 nucleosomes in the absence of CENP-A^Cnp1^ may be critical in earmarking these chromosomal regions for CENP-A^Cnp1^ assembly.

The analyses presented here suggest that the previously described H3 placeholder nucleosomes deposited within the central domain during S phase (Shukla et al., 2017) contain H2A.Z^Pht1^ and H3K56ac. The events that subsequently lead to the replacement of placeholder H3 nucleosomes with CENP-A^Cnp1^ nucleosomes are largely unknown. Transcription of centromere DNA has been widely implicated in CENP-A^Cnp1^ deposition in several systems (Duda et al., 2017; Rosic and Erhardt, 2016). RNAPII transcription is an effective chromatin remodeler, thus removal of H2A.Z^Pht1^ containing H3 nucleosomes deposited at *S. pombe* centromeres in S phase may occur as a result of promoter-driven nucleosome disassembly allowing the assembly of CENP-A^Cnp1^ nucleosomes in their place. In agreement with this possibility, elevated elongating RNAPII levels have been shown to accumulate on central domain DNA during G2 coincident with H3 eviction (Shukla et al., 2017). In addition, the Ino80 remodeling complex was recently found to be recruited to CENP-A^Cnp1^ chromatin in *S. pombe*, and required for the removal of histone H3 from within centromeres (Choi et al., 2017). Ino80 has been reported to remove H2A.Z from nucleosomes (Watanabe and Peterson, 2010), however it remains to be determined if this activity contributes to the conversion of S phase H3 placeholder nucleosomes to CENP-A nucleosomes.

A key feature of our model is that the inclusion of H2A.Z^Pht1^ in placeholder H3 nucleosomes at centromeres drives their destabilization, disassembly and subsequent replacement by CENP-A^Cnp1^ nucleosomes. Nucleosome core particles assembled *in vitro* with the vertebrate H3.3 variant and H2A.Z are known to be less stable on DNA than those assembled with canonical H3.1 and H2A.Z (Jin and Felsenfeld, 2007). In *S. pombe* all histone H3 is more closely related to vertebrate H3.3 (Choi et al., 2005; Hake and Allis, 2006), thus *S. pombe* nucleosomes containing H2A.Z^Pht1^ are likely to be innately unstable. Interestingly, in *S.cerevisiae*, *S. pombe* and human cells endogenous or overexpressed CENP-A is detected at regions of high nucleosome turnover (i.e. the 5’ ends of genes), or where H3.3 and/or H2A.Z is incorporated (Choi et al., 2011; Hildebrand and Biggins, 2016; Lacoste et al., 2014). Such observations generally support the idea that CENP-A is preferentially deposited at sites where H3 is actively evicted.

H2A.Z resides in centromeric regions on mammalian chromosomes where it contributes to centromere structure and function (Foltz et al., 2006; Greaves et al., 2007). Defective chromosome segregation is observed in several eukaryotes with aberrant H2A.Z deposition and has been attributed to impaired centromeric cohesion resulting from improper heterochromatin formation (Boyarchuk et al., 2011). Notably, loss of *S. pombe* H2A.Z^Pht1^ or Swr1C-associated Msc1 is synthetically lethal with mutations in CENP-A^Cnp1^ and its loading factors and Msc1 overexpression rescues these lethal mutations (Ahmed et al., 2007; Gao et al., 2017; Kim et al., 2009). Such genetic interactions underscore a role for H2A.Z^Pht1^ in CENP-A^Cnp1^ deposition, kinetochore integrity and chromosome segregation. It remains to be determined whether H2A.Z at mammalian centromeres, and factors such as SRCAP^Swr1^, which mediate mammalian H2A/H2B-to-H2A.Z/H2B dimer exchange (Ruhl et al., 2006; Wong et al., 2007), influence CENP-A deposition and kinetochore integrity in mammals. The fact that H3.3 is deposited as a placeholder in S phase (Dunleavy et al., 2011) and that H2A.Z levels increase during mitosis (Nekrasov et al., 2012), prior to CENP-A replenishment at mammalian centromeres in G1 (Dunleavy et al., 2011; Jansen et al., 2007), predict that H2A.Z may indeed be involved.

## AUTHOR CONTRIBUTIONS

Conceptualization, R.K-S. and R.C.A.; Investigation, R. K-S., and L.S.; Formal analysis, A.R.W.K.; Resources, A.R.W.K., C.S. and J.R.; Writing – Original Draft, R.K-S. and R. C.A.; Writing – Review & Editing, R.K-S., R.C.A., L.S., A.R.W.K. and C.S.; Funding Acquisition, R.C.A.

## ACKNOWLEDGEMENTS

We thank Mitsuhiro Yanagida, Vincent Vanoosthuyse, Karl Ekwall and Songtao Jia for sharing strains and Takeshi Urano for providing the H3K9me2 antibody (mAb5.1.1). We thank Alison Pidoux for comments on the manuscript, and Flavia de Lima Alves and Pin Tong for mass spectrometry and bioinformatics support respectively.

R. K-S. was partly supported by Peter und Traudl Postdoctoral Fellowship. L.S. was supported in part by EC FP7 Marie Curie International Incoming (PIIF-GA-2010-275280) and EMBO Long Term Fellowships (ALTF 1491-2010). J.R. is a Wellcome Senior Research Fellow (103139) and holds a Wellcome instrument grant (108504). R.C.A. is a Wellcome Principal Research Fellow (200885); core support for the Wellcome Centre for Cell Biology (203149).

## METHOD DETAILS

### Strains, Plasmids

*S. pombe* strains used in this study are listed in Table S2. Standard methods were used for fission yeast growth, genetics and manipulation. PCR-based methods (Bahler et al., 1998) were used for gene deletions and epitope tagging of proteins. Centromeric plasmid transformations were carried out by electroporation. Plasmid sequences and detailed maps are available upon request. Five-fold serial dilutions of the indicated strains were spotted onto YES and PMG media with/without arginine for silencing assays.

### Immunoaffinity purification and mass spectrometry

For CENP-A^Cnp1^, 5 g of pulverized *S. pombe* cells expressing GFP-tagged CENP-A^Cnp1^ were used for immunoprecipitation with anti-GFP antibody A11122 (Life Technologies) coupled to Protein G Dynabeads (Life Technologies), alongside an untagged control. After washes, Dynabeads with immunoprecipitated material were subjected to on-bead tryptic digestion following which the samples were treated as described (Subramanian et al., 2014) and prepared for LC-MS/MS analysis. LC-MS-analyses were performed on a Q Exactive mass spectrometer (Thermo Fisher Scientific) coupled, on-line, to an Ultimate 3000 RSLCnano Systems (Dionex, Thermo Fisher Scientific). Peptides were separated on a 50 cm EASY-Spray column (Thermo Scientific) assembled in an EASY-Spray source (Thermo Scientific) and operated at 50°C. Mobile phase A consisted of 0.1% formic acid in water while mobile phase B consisted of 80% acetonitrile and 0.1% formic acid. Peptides were loaded onto the column at a flow rate of 0.3 µL min^−1^ and eluted at a flow rate of 0.2 µL min^−1^ according to the following gradient: 2 to 40% buffer B in 90 min, then to 95% in 11 min and 2 to 40% in 120 min and then to 95% in 11 min. The resolution for the FTMS spectra was set at 70,000 (scan range 350-1400 m/z) and the ten most intense peaks with charge ≥ 2 of the MS scan were selected with an isolation window of 2.0 Thomson for MS2 (filling 1.0E6 ions for MS scan, 5.0E4 ions for MS2, maximum fill time 60 ms, dynamic exclusion for 50 s). The MaxQuant software platform (Cox and Mann, 2008) version 1.5.2.8 was used to process raw files and search was conducted against *Schizosaccharomyces pombe* complete proteome set of PomBase (released in March, 2016), using Andromeda peptide search engine (Cox et al., 2011). The first search peptide tolerance was set to 20 ppm while the main search peptide tolerance was set to 4.5 ppm. Isotope mass tolerance was set to 2 ppm and maximum charge was set to 7. Maximum of two missed cleavages were allowed. Carbamidomethylation of cysteine was chosen as fixed modification. Oxidation of methionine and acetylation of the N-terminal were chosen as variable modifications. The average number of unique peptides corresponding to proteins that were reproducibly enriched in the epitope-tagged samples is presented.

### Co-immunoprecipitation and Western analyses

For co-immunoprecipitation experiments, pulverized *S. pombe* cells expressing the indicated epitope-tagged proteins were used for immunoprecipitation using anti-GFP antibody A11122 (Life Technologies) coupled to Protein G Dynabeads (Life Technologies). The anti-Myc antibody 9B11 (Cell Signaling) or anti-GFP antibody (Roche) was used for Western analyses as indicated.

### Centromeric plasmid selection system and stability

Plasmids bearing only the central domain of centromere 2 sequence (pCC2) or central domain with flanking heterochromatin (pHCC2) have been described earlier (Folco et al., 2008). Both pCC2 & pHCC2 plasmids carry *ura4*^+^ and *sup3-e* (suppressor of *ade6-704*) and *kan*R selection systems in addition to a minimal ars1 element to ensure efficient replication in *S. pombe*. In all strains tested, including wild type, the central domain of endogenous centromere 2 was replaced with the central domain of centromere 1 (*cc2*∆::*cc1*) (Catania et al., 2015) so that the only copy of central domain of centromere 2 sequence (*cnt2*) is carried by the minichromosome. Cells without *ura4*^+^ cannot grow on minus-uracil plates, while *ade6-704* cells do not grow without adenine and form red colonies on 1/10th adenine plates. The *sup3-e* tRNA gene suppresses a premature stop in *ade6-704*, allowing growth on minus-adenine plates. The acentric pcc2 plasmid is mitotically unstable and rapidly lost and therefore forms red colonies on 1/10th adenine indicator plates but is maintained in –ura – ade selective media. Cells containing pHCC2 form a high percentage of white or sectored colonies on 1/10th adenine indicator plates, demonstrating their relative mitotic stability in wild-type cells. In cells lacking Clr4, however, their mitotic stability is lost due to a lack of heterochromatin-dependent centromeric cohesion and inability to assemble CENP-A. To confirm that plasmids were behaving episomally and had not integrated, a plasmid stability test was performed at the time of fixation for ChIP. Cells (100 – 1000) were plated onto YES supplemented with 1/10th adenine and allowed to form colonies. Samples exhibiting no integrations were used for ChIP.

### Chromatin Immunoprecipitation (ChIP)

The indicated *S. pombe* strains were grown in YES media at 32° C. For cell cycle experiments, fission yeast cells carrying the *cdc25-22* mutation (Moreno et al., 1989) were grown overnight in YES media at 25° C. Exponentially growing cells were shifted to 36° C for 3.5 hours, cooled to 25° C, and further cultured to release them synchronously into the cell 3.6 cycle. Cells were then harvested and fixed every 15 minutes for the first 120 minutes and at 30-minute intervals thereafter. Septation index was monitored to ensure comparable synchronization of cultures among all cell cycle experiments. For ChIPs on centromeric plasmids, cells harboring pCC2 or pHCC2 were grown in PMG media lacking adenine and uracil, at 32° C. To confirm that plasmids were behaving episomally and had not integrated, a plasmid stability test was performed at the time of fixation. Cells (100–1000) were plated onto YES supplemented with 1/10th adenine and allowed to form colonies. Samples exhibiting no integrations were used for ChIP.

ChIP was performed as described (Keller et al., 2013), with the following modifications. Cells were fixed with 1% formaldehyde (Sigma) for 15 min at room temperature and lysed using a Mini Beadbeater (Biospec Products). Cell lysates were sonicated in a Bioruptor (Diagenode) (30/40 min, 30 s On and 30 s Off at ‘High’ (200W) position). For CENP-A^Cnp1^ ChIPs, anti-CENP-A^Cnp1^ serum was used with Protein G sepharose beads (GE Healthcare). For all other ChIPs, Protein G Dynabeads (Life Technologies) were used along with anti-H3K9me2 (mAb5.1.1), anti-H3K56ac (ActivMotif), anti-Myc 9B11 (Cell Signaling), anti-HA 12CA5 (Roche) or anti-Flag M2 antibody (Sigma), as appropriate. Immunoprecipitated DNA was recovered using Qiagen Min-Elute PCR purification kit. ChIPs were analyzed by real-time PCR using Lightcycler 480 SYBR Green (Roche) with primers specific to the central cores of centromere 2 (*cnt2*) or centromeres 1/3 (*cnt1*), or *act1*. ChIP enrichments were calculated as % DNA immunoprecipitated at the locus of interest relative to the corresponding input samples, relative to H3 or % DNA immunoprecipitated at the *act1* locus and/or normalized to *cnt1* or wild type. For cell cycle experiments data points were normalized to T0 time point i.e., before cells were released following arrest at 36° C. Data from 2 representative experiments are presented in each figure.

### ChIP-sequencing

ChIP-seq libraries were generated using an Illumina-based protocol with custom reagents using NEXTFlex-96 DNA Barcodes (Bioo Scientific). Briefly, immunoprecipitated DNA was end-repaired using a combination of T4 DNA polymerase, *E. coli* DNA Pol I large fragment (Klenow polymerase) and T4 polynucleotide kinase. The blunt phosphorylated ends were treated with Klenow fragment (3’ to 5’ exo minus) and dATP to yield a protruding 3-’A’ base for ligation of Illumina adapters, which have a single ’T’ base overhang at the 3’ end. After adapter ligation, DNA fragments were size-selected using AmpureXP beads to remove excess adapters and adapter-modified DNA fragments were enriched by PCR amplification with Illumina primers for 12 cycles. The purified ChIP-seq libraries thus generated were validated and subsequently captured on an Illumina flow cell for cluster generation. Libraries were sequenced on the Illumina HiSeq2500 or 4000 system following the manufacturer’s protocols.

BWA (Li and Durbin, 2009, 2010) or BOWTIE2 (Langmead and Salzberg, 2012) was used to map ChIP-seq reads onto the *S. pombe* genome assembly EF2 (ASM294v2) or modified genome assembly generated based on ASM294v2, where centromere 2 was replaced with sequence from centromere 1 to mimic (cc2∆::cc1) and to include pHCC2 or pCC2 plasmid sequence. Indexed, sorted bam files were created for each dataset using SAMtools (Li et al., 2009). The program bamCoverage from the deepTools suite version 2.5.3 (Ramirez et al., 2016) or BEDtools (Quinlan and Hall, 2010) was used to create bigwig files from the mapped bam files and scaled using reads per kilobase of transcript per million mapped reads [RPKM], and SeqPlots (Stempor and Ahringer, 2016) was used to plot the occupancy over transcription start sites [TSS] as annotated in PomBase. To compare multiple ChIP-seq experiment traces, each trace was plotted on axes of the same scale: the y-axes are plotted in RPKM, and the axis scale for each trace is noted in the figures.

### Quantification And Statistical Analysis

Statistical parameters are reported in the figure legends.

## Data And Software Availability

The accession number for ChIP-seq datasets reported in this paper is GEO: GSE106585. The accession number for the mass spectrometry raw data is ProteomeXchange: PXD008126. Original datasets have been deposited to Mendeley Data: http://dx.doi.org/10.17632/52w9sp2k6p.1.

## SUPPLEMENTARY FIGURE LEGENDS

**Figure S1:**
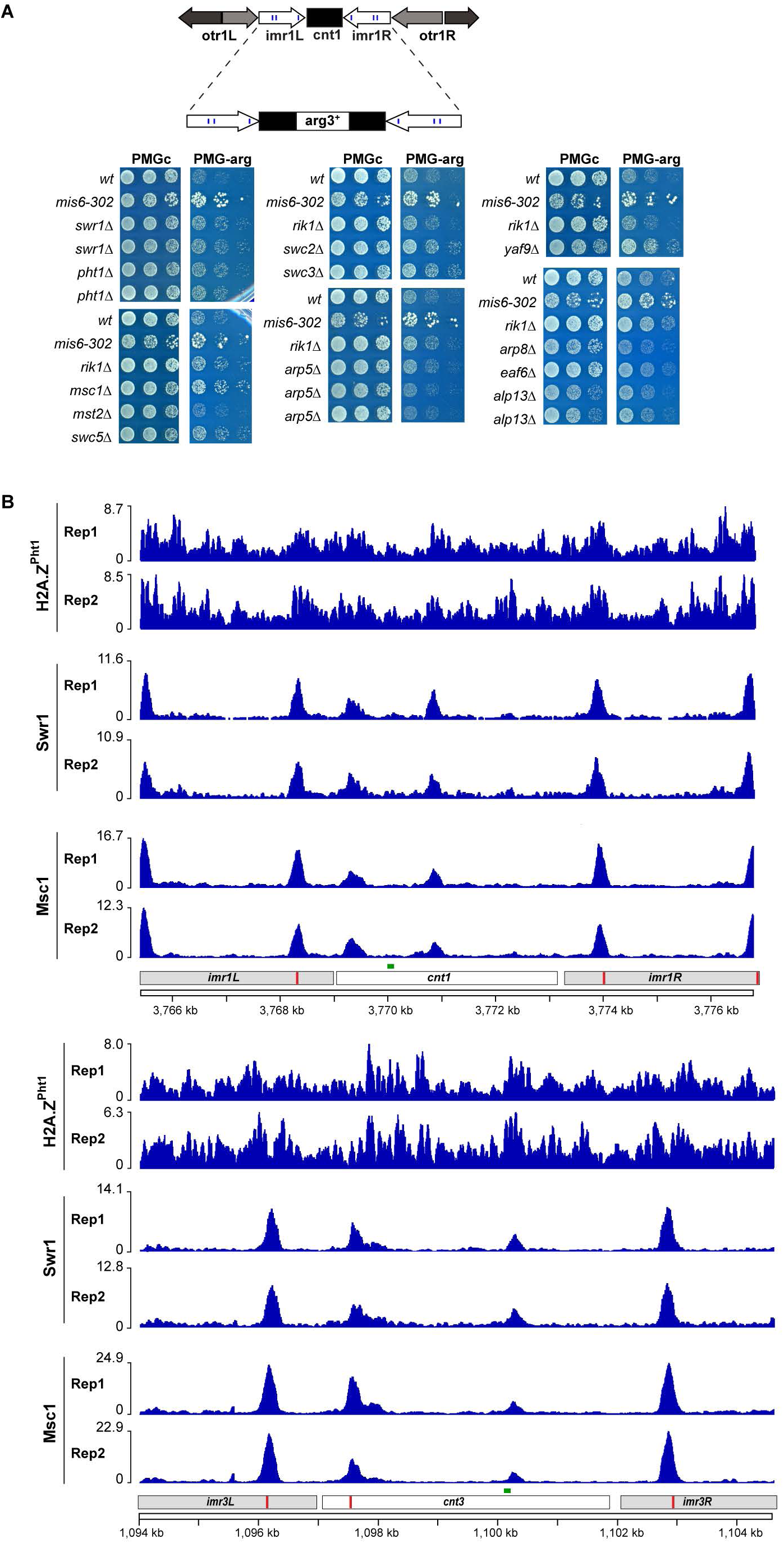
(related to Figure 1) (A) Centromere marker silencing assay showing the effect of deleting Swr1, Ino80, Mst1 complex components or Mst2 on silencing of the *arg3* reporter gene inserted in the central domain of centromere 1. (B) ChIP-seq profile for H2A.Z^Pht1^, Swr1 and Msc1 across central domains of centromeres 1 and 3. Chromosome positions (kb) and annotation indicated. Scale represents normalized reads in RPKM.

**Figure S2:**
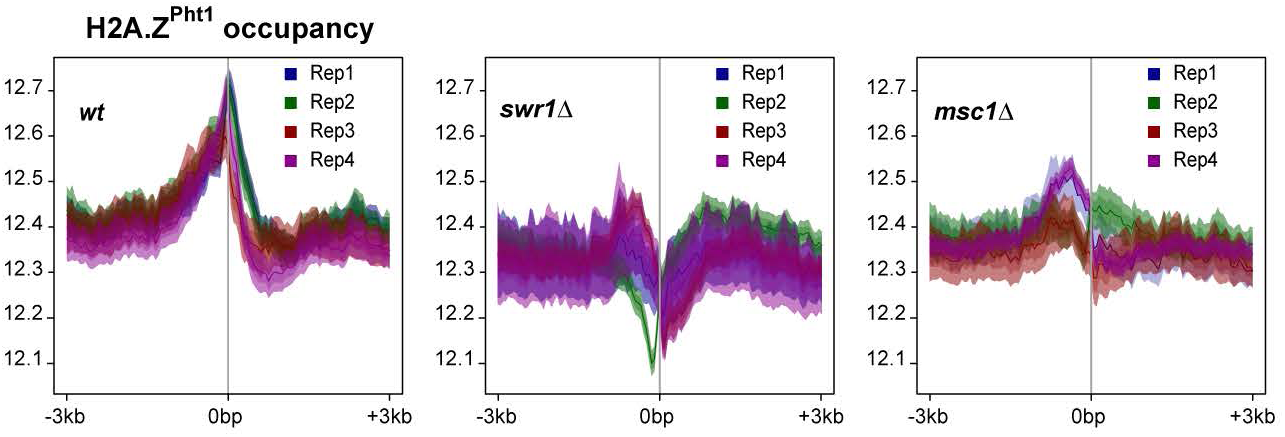
(related to Figure 2) (A) TSS profile showing changes in nucleosome occupancy at TSS (± 3kb) in *swr1∆* and *msc1∆* compared to *wt*. Scale represents normalized reads in RPKM.

**Figure S3:**
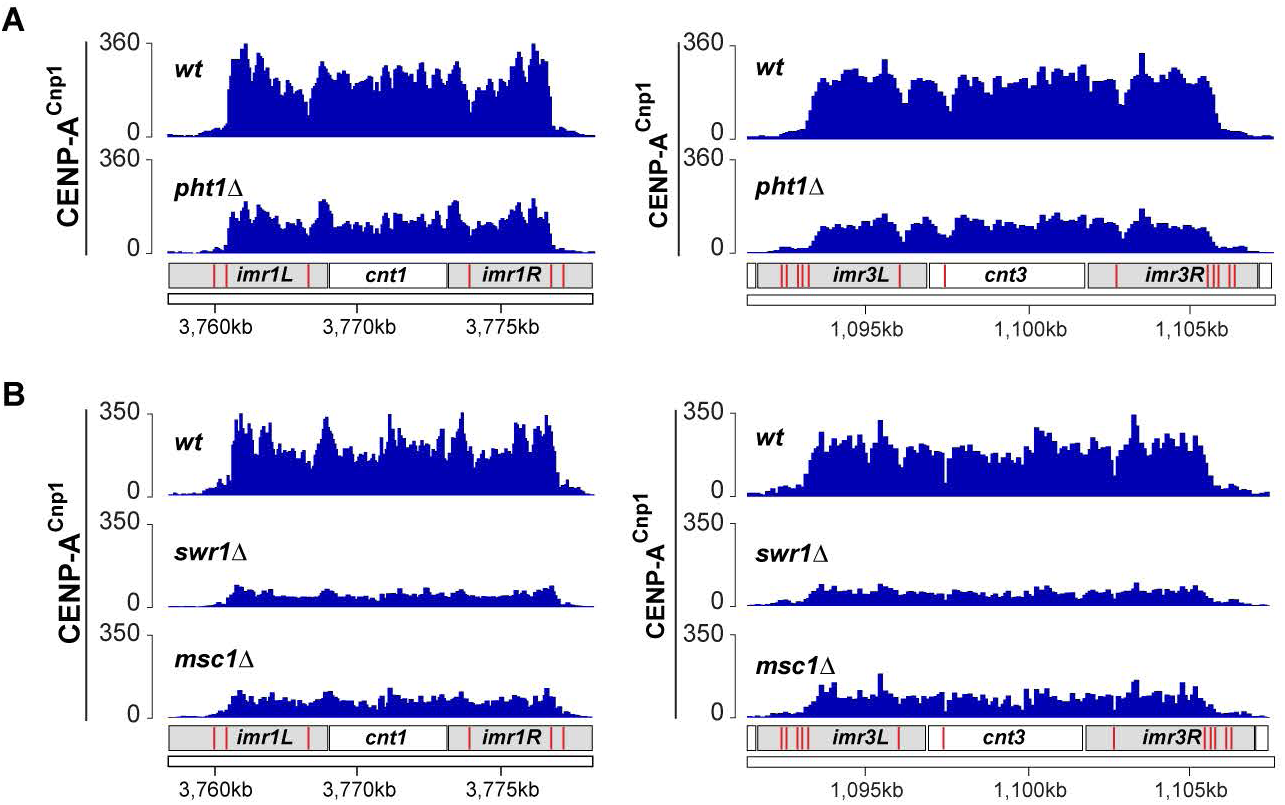
(related to Figure 3) (A) CENP-A^Cnp1^ ChIP-seq profile across centromeres 1 and 3 in *wt*, and *pht1∆*. Chromosome positions (kb) and annotation indicated. Scale represents normalized reads in RPKM. (B) CENP-A^Cnp1^ ChIP-seq profile across centromeres 1 and 3 in *wt*, *swr1∆* and *msc1∆.* Chromosome positions (kb) and annotation indicated. Scale represents normalized reads in RPKM.

**Figure S4:**
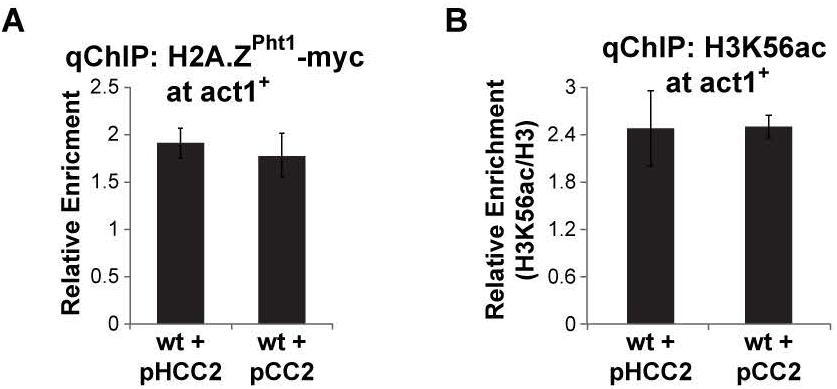
(related to Figure 4) (A) qChIP analysis showing H2A.Z^Pht1^ levels at *act1*^+^ in *wt* strains harboring pCC2 or pHCC2 minichromosomes. Mean ± SEM of n≥3 independent experiments is shown. (B) qChIP analysis showing H3K56ac levels at *act1*^+^ in *wt* strains harboring pCC2 or pHCC2 minichromosomes. Mean ± SEM of n≥3 independent experiments is shown.

**Figure S5:**
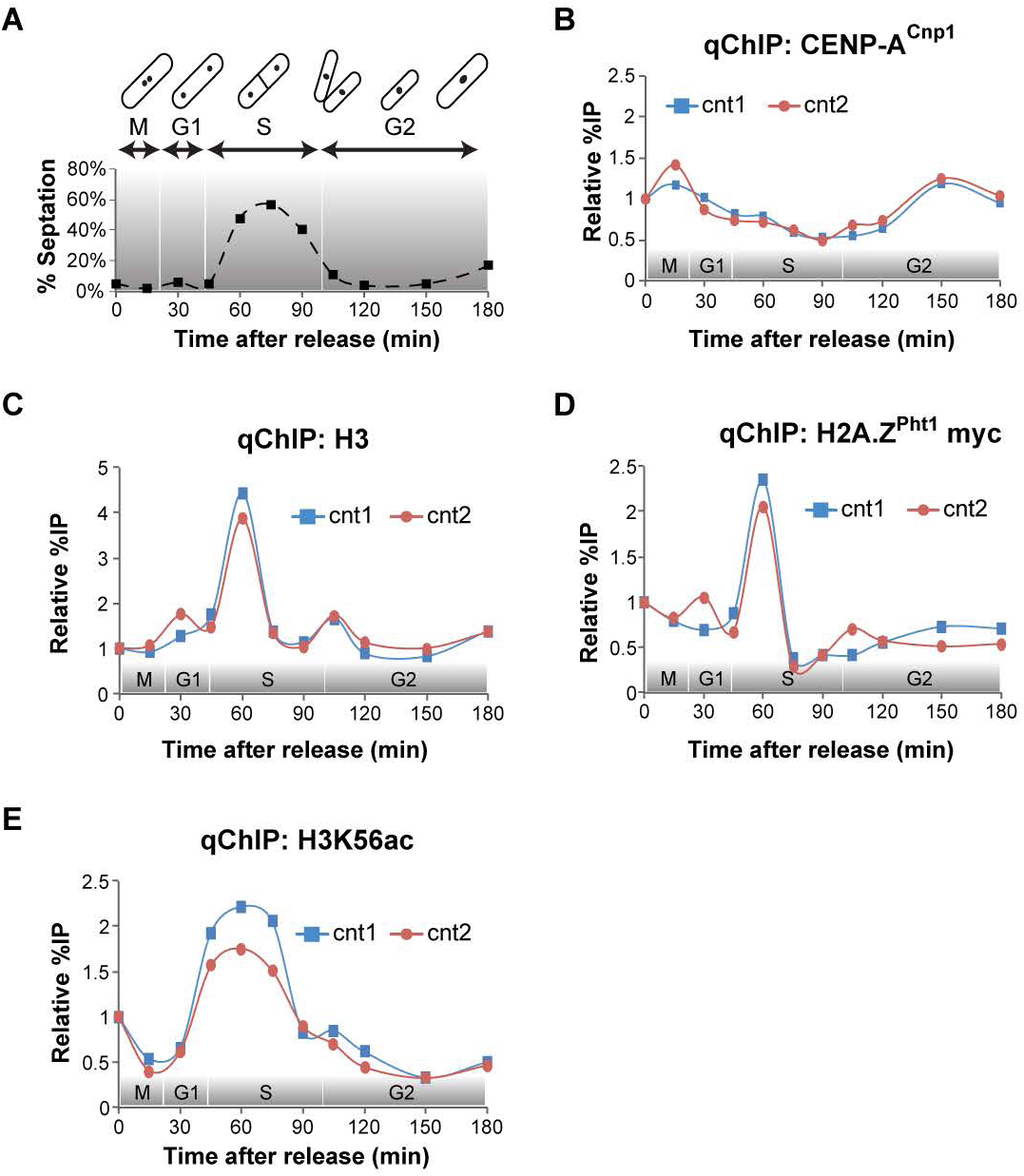
(related to Figure 5) (A) Septation index plot to monitor cell cycle synchrony. Expected cell cycle phases shown above. (B) Independent replicate of qChIP for CENP-A^Cnp1^ at *cnt1* and *cnt2* in cell cycle synchronized cells. (C) Independent replicate of qChIP for histone H3 at *cnt1* and *cnt2* in cell cycle synchronized cells. (D) Independent replicate of qChIP for H2A.Z^Pht1^ at *cnt1* and *cnt2* in cell cycle synchronized cells. (E) Independent replicate of qChIP for H3K56ac at *cnt1* and *cnt2* in cell cycle synchronized cells.

**Figure S6:**
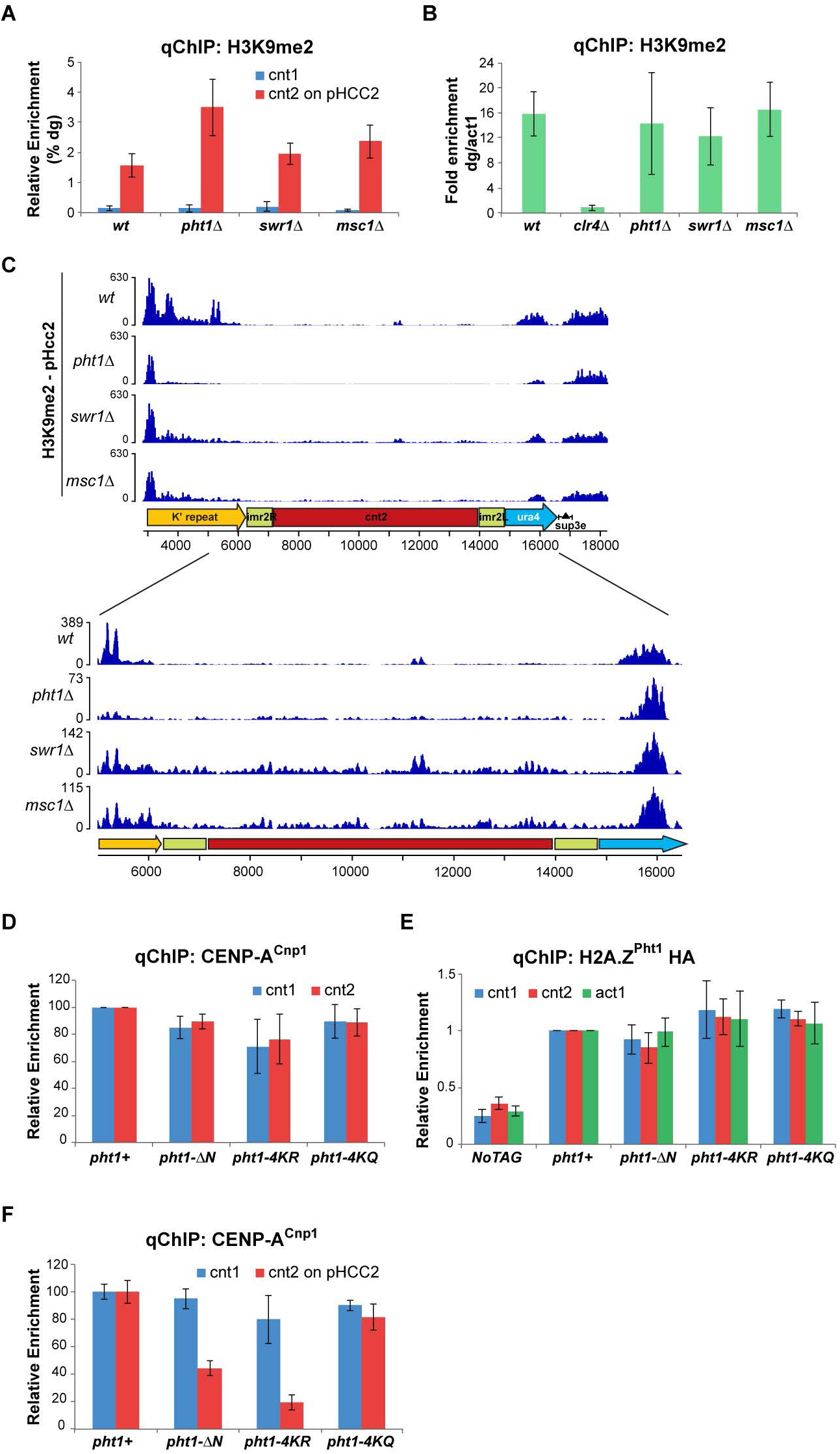
(related to Figure 6) (A) qChIP analysis showing H3K9me2 levels at endogenous *cnt1*, and *cnt2* on pHCC2 relative to *dg*. Mean ± SEM of n=3 independent experiments is shown. (B) qChIP analysis showing H3K9me2 levels at endogenous *dg* relative to *act1* in strains harboring pHCC2 minichromosome. Mean ± SEM of n=3 independent experiments is shown. (C) Upper panel shows H3K9me2 ChIP-seq profile across pHCC2 minichromosome. Lower panel shows zoomed-in profile across central domain DNA sequence on pHCC2. Minichromosome coordinates (kb) and map of pHCC2 are shown. Scale represents normalized reads in RPKM. (D) qChIP analysis showing CENP-A^Cnp1^ levels at endogenous *cnt1* and *cnt2* in H2A.Z^Pht1^ acetylation mutants normalized to *wt* (*pht1+*). Mean ± SEM of n=3 independent experiments is shown. (E) qChIP analysis showing H2A.Z^Pht1^ levels normalized to *wt* (*pht1+*) in H2A.Z^Pht1^ acetylation mutants. Mean ± SEM of n=3 independent experiments is shown. (F) qChIP analysis showing H2A.Z^Pht1^ levels at endogenous *cnt1*, and *cnt2* on pHCC2 normalized to *wt* (*pht1+*). Mean ± SEM of n=3 independent experiments is shown.

**Table S1.**
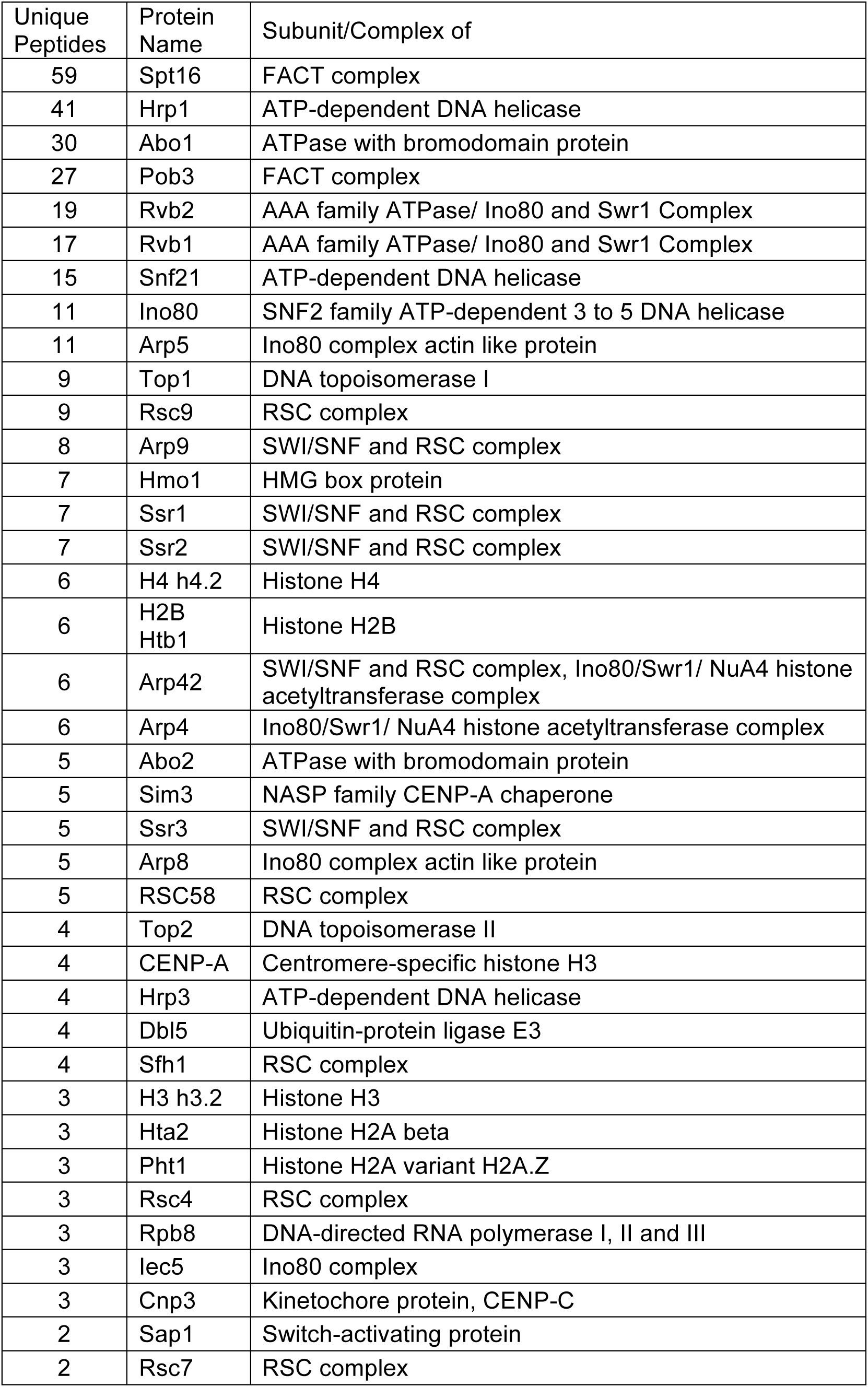
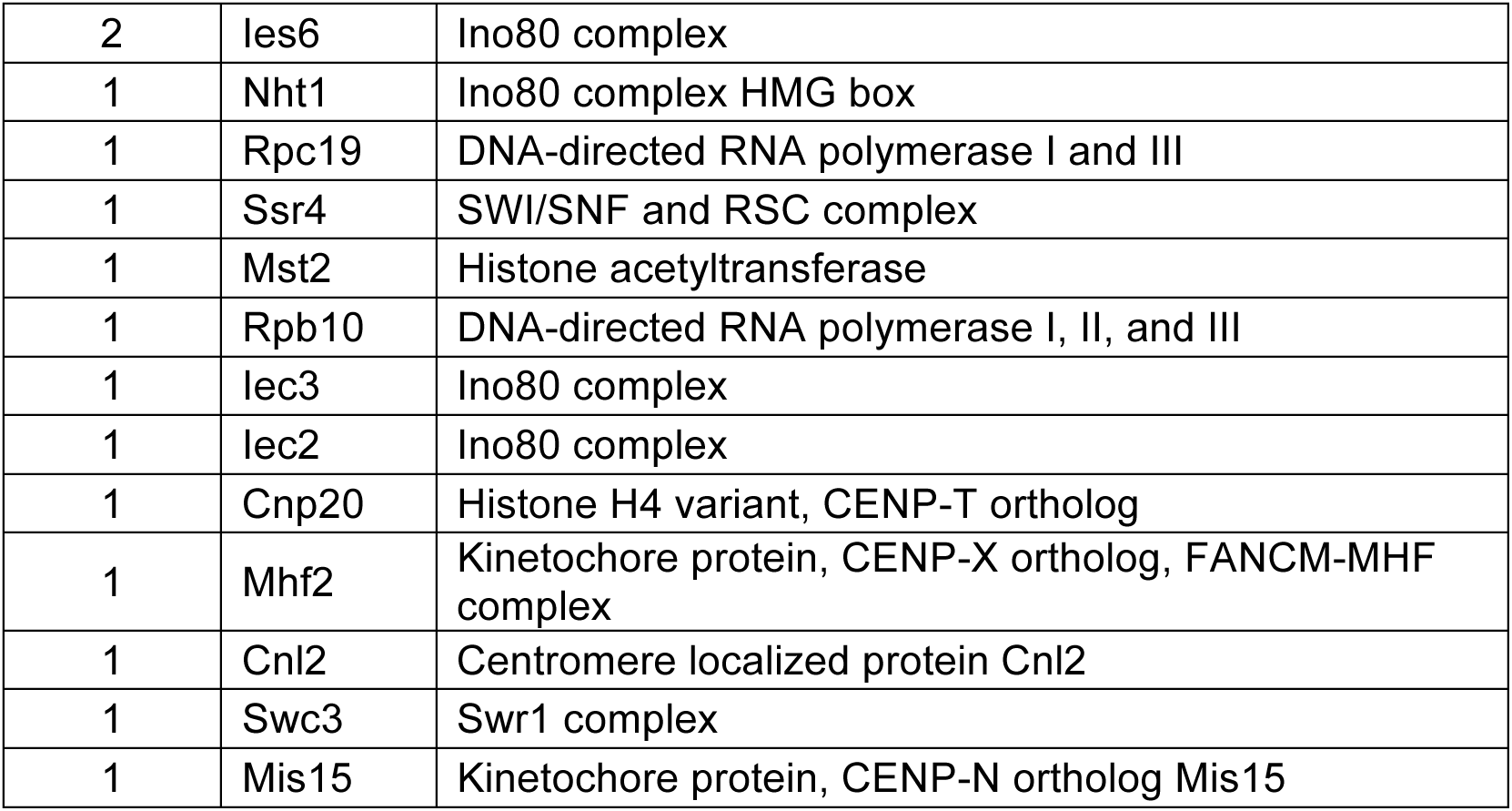
(Related to Figure 1): List of factors detected by LC MS/MS analysis of GFPCENP-A^Cnp1^

**Table S2.** (Related to STAR Methods): List of yeast strains used in this study

| Name | Genotype | Ref |
| --- | --- | --- |
| FY2221 | h- TM1(Ncol):arg3 ade6-210/D1 D4 his3-D1 leu1-32 ura4-D18 | Allshire Lab Collection |
| FY3027 | h+ ade6-210 D3 his3-D1 leu1-32 ura4-D18/DS-E TM1:srg3 TM3:ade6 tel1L:his3 otr2:ura4 | Allshire Lab Collection |
| FY3606 | h? rik1::leu2, TM1:arg3 TM3:ade6, otr2:ura4, tel1L:his3, ade6-210, 4, his3D1, leu1-32, ura4D18 | Allshire Lab Collection |
| FY5691 | h mis6-302 cnt1:arg3 cnt3:ade6 otr2:ura4 tel1L:his3 ade6-210 leu1-32 ura4-D18 D4 his3-D1 | Allshire Lab Collection |
| A2549 | h- leu1-32 ura4-D18 lys1 GFP-cnp1-NAT | Takayama et. al., 2008 |
| A7373 | h- leu1-32 ura4-D18/DS-E ade6-704:hphMX6 cc2Δ6kb:cc1 | Catania et al., 2015 |
| A7377 | h- leu1-32 ura4-D18/DS-E ade6-704:hphMX6 cc2Δ6kb:cc1 clr4Δ::NatMX | Catania et al., 2015 |
| B0659 | h+ ade6-M216 ura4-D18 leu1-32 his3-D1 pht1.HA3-KanMX | Kim et al., 2009 |
| B0660 | h+ ade6-M216 ura4-D18 leu1-32 his3-D1 pht1-ΔN.HA3-KanMX | Kim et al., 2009 |
| B0663 | h+ ade6-M216 ura4-D18 leu1-32 his3-D1 pht1-4KR.HA3-KanMX | Kim et al., 2009 |
| B0664 | h+ ade6-M216 ura4-D18 leu1-32 his3-D1 pht1-4KQ.HA3-KanMX | Kim et al., 2009 |
| B2148 | h- leu1-32 ura4-D18/DS-E ade6-704:hphMX cc2Δ6kb:cc1 swr1Δ::NatMX | This Study |
| B2149 | h- TM1(Ncol):arg3 ade6-210/D1 D4 his3-D1 leu1-32 ura4-D18 swr1Δ::NatMX | This Study |
| B2150 | h- TM1(Ncol):arg3 ade6-210/D1 D4 his3-D1 leu1-32 ura4-D18 swr1Δ::NatMX | This Study |
| B2154 | h- leu1-32 ura4-D18/DS-E ade6-704:hphMX cc2Δ6kb:cc1 pht1Δ::NatMX | This Study |
| B2157 | h- TM1(Ncol):arg3 ade6-210/D1 D4 his3-D1 leu1-32 ura4-D18 pht1Δ::NatMX | This Study |
| B2158 | h- TM1(Ncol):arg3 ade6-210/D1 D4 his3-D1 leu1-32 ura4-D18 pht1Δ::NatMX | This Study |
| B2196 | h- leu1-32 ura4-D18 lys1 GFP-cnp1-NAT msc1-Flag::KanMX | This Study |
| B2200 | h- leu1-32 ura4-D18 lys1 GFP-cnp1-NAT swr1-myc::KanMX | This Study |
| B2202 | h- leu1-32 ura4-D18 lys1 GFP-cnp1-NAT pht1-myc::KanMX | This Study |
| B2219 | h- leu1-32 ura4-D18 lys1 GFP-cnp1-NAT pht1Δ::hphMX | This Study |
| B2222 | h- leu1-32 ura4-D18 lys1 GFP-cnp1-NAT swr1Δ::hphMX | This Study |
| B2223 | h- leu1-32 ura4-D18/DS-E ade6-704:hphMX cc2Δ6kb:cc1 msc1Δ::NatMX | This Study |
| B2224 | h- leu1-32 ura4-D18 lys1 GFP-cnp1-NAT msc1Δ::hphMX | This Study |
| B2335 | h? leu1-32 ura4-D18 lys1 cdc25-22 GFP-cnp1-NAT pht1-myc::KanMX | This Study |
| B2408 | h+ ade6-210 D3 his3-D1 leu1-32 ura4-D18/DS-E<br>TM1:arg3 TM3:ade6 tel:his3 otr2:ura4 swc2Δ::NatMX | This Study |
| B2409 | h+ ade6-210 D3 his3-D1 leu1-32 ura4-D18/DS-E<br>TM1:arg3 TM3:ade6 tel:his3 otr2:ura4 swc3Δ::NatMX | This Study |
| B2413 | h+ ade6-210 D3 his3-D1 leu1-32 ura4-D18/DS-E<br>TM1:arg3 TM3:ade6 tel:his3 otr2:ura4 yaf9Δ::NatMX | This Study |
| B2419 | h+ ade6-210 D3 his3-D1 leu1-32 ura4-D18/DS-E<br>TM1:arg3 TM3:ade6 tel:his3 otr2:ura4 arp5Δ::NatMX | This Study |
| B2420 | h+ ade6-210 D3 his3-D1 leu1-32 ura4-D18/DS-E<br>TM1:arg3 TM3:ade6 tel:his3 otr2:ura4 arp5Δ::NatMX | This Study |
| B2421 | h+ ade6-210 D3 his3-D1 leu1-32 ura4-D18/DS-E<br>TM1:arg3 TM3:ade6 tel:his3 otr2:ura4 arp5Δ::NatMX | This Study |
| B2423 | h+ ade6-210 D3 his3-D1 leu1-32 ura4-D18/DS-E<br>TM1:arg3 TM3:ade6 tel:his3 otr2:ura4 arp8Δ::NatMX | This Study |
| B2424 | h+ ade6-210 D3 his3-D1 leu1-32 ura4-D18/DS-E<br>TM1:arg3 TM3:ade6 tel:his3 otr2:ura4 eaf6Δ::NatMX | This Study |
| B2425 | h+ ade6-210 D3 his3-D1 leu1-32 ura4-D18/DS-E<br>TM1:arg3 TM3:ade6 tel:his3 otr2:ura4 alp13Δ::NatMX | This Study |
| B2426 | h+ ade6-210 D3 his3-D1 leu1-32 ura4-D18/DS-E<br>TM1:arg3 TM3:ade6 tel:his3 otr2:ura4 alp13Δ::NatMX | This Study |
| B2433 | h? leu1-32 ura4-D18 lys1 cdc25-22 GFP-cnp1-NAT<br>pht1-myc::KanMX swr1Δ::hphMX | This Study |
| B2437 | h? leu1-32 ura4-D18 lys1 cdc25-22 GFP-cnp1-NAT<br>pht1-myc::KanMX msc1Δ::hphMX | This Study |
| B2538 | h+ ade6-210 D3 his3-D1 leu1-32 ura4-D18/DS-E<br>TM1:arg3 TM3:ade6 tel:his3 otr2:ura4 eaf7Δ::NatMX | This Study |
| B2539 | h+ ade6-210 D3 his3-D1 leu1-32 ura4-D18/DS-E<br>TM1:arg3 TM3:ade6 tel:his3 otr2:ura4 eaf7Δ::NatMX | This Study |
| B2555 | h+ ade6-210 D3 his3-D1 leu1-32 ura4-D18/DS-E<br>TM1:arg3 TM3:ade6 tel:his3 otr2:ura4 msc1Δ::hphMX | This Study |
| B2561 | h? leu1-32 ura4-D18/DS-E ade6-704:hphMX<br>cc2Δ6kb:cc1 pht1-ΔN.HA3-KanMX | This Study |
| B2563 | h? leu1-32 ura4-D18/DS-E ade6-704:hphMX<br>cc2Δ6kb:cc1 pht1-4KR.HA3-KanMX | This Study |
| B2565 | h? leu1-32 ura4-D18/DS-E ade6-704:hphMX<br>cc2Δ6kb:cc1 pht1-4KQ.HA3-KanMX | This Study |
| B2583 | h- leu1-32 ura4-D18/DS-E ade6-704:hphMX<br>cc2Δ6kb:cc1 pht1-myc::KanMX | This Study |
| B2602 | h+ ade6-210 D3 his3-D1 leu1-32 ura4-D18/DS-E<br>TM1:arg3 TM3:ade6 tel:his3 otr2:ura4 mst2Δ::hphMX | This Study |
| B2610 | h+ ade6-210 D3 his3-D1 leu1-32 ura4-D18/DS-E<br>TM1:arg3 TM3:ade6 tel:his3 otr2:ura4 swc5Δ::NatMX | This Study |

